# KMT1/Suv39 methyltransferase family regulates peripheral heterochromatin tethering via histone and non-histone protein methylations

**DOI:** 10.1101/240952

**Authors:** Radhika Arasala Rao, Alhad Ashok Ketkar, Neelam Kedia, Vignesh K Krishnamoorthy, Vairavan Lakshmanan, Pankaj Kumar, Abhishek Mohanty, Shilpa Dilip Kumar, Sufi O Raja, Akash Gulyani, ChandraPrakash Chaturvedi, Marjorie Brand, Dasaradhi Palakodeti, Shravanti Rampalli

## Abstract

Euchromatic histone methyltransferases (EHMTs), members of the KMT1 family, methylate histone and non-histone proteins. Here we uncover a novel role for EHMTs in regulating heterochromatin anchorage to the nuclear periphery (NP) via non-histone (LaminB1) methylations. We show that EHMTs methylates and stabilizes LaminB1 (LMNB1), which associates with the H3K9me2-marked peripheral heterochromatin. Loss of LMNB1 methylation or EHMTs abrogates the heterochromatin anchorage from the NP. We further demonstrate that the loss of EHMTs induces many hallmarks of aging including global reduction of H3K27methyl marks along with altered nuclear-morphology. Consistent with this, we observed a gradual depletion of EHMTs, which correlates with loss of methylated LMNB1 and peripheral heterochromatin in aging human fibroblasts. Restoration of EHMT expression reverts peripheral heterochromatin defect in aged cells. Collectively our work elucidates a new mechanism by which EHMTs regulate heterochromatin domain organization and reveals their impact on fundamental changes associated with the intrinsic aging process.

## Introduction

The Euchromatic histone lysine methyltransferases G9a, encoded by EHMT2, and GLP, encoded by EHMT1 (KMT1/Suv39 methyltransferase family), are present as heteromeric complex and negatively regulate gene transcription. The SET domain of EHMT catalyzes mono and dimethylation of lysine residues at histone3 (H3) *in vitro* and *in vivo* [1–5]. H3K9me2 deposited by EHMT1/2 complex demarcates heterochromatin, particularly non-genic regions and is prevalent in gene deserts, pericentromeric and subtelomeric regions, with little being observed at individual active or silent genes. Non-coding and gene containing DNA present at the NP are also marked by the presence of H3K9me2 which spans several megabases [6,7]. Specifically, these domains are strongly correlated with binding of LMNB1 and are depleted of H3K4me3 and RNA Polymerase II activity [6]. These data suggest that H3K9me2 domains are critical determinants of higher-order chromosome structure in association with the nuclear lamina (NL).

In mammalian cells, the NL acts as a hub for multiple cellular functions including chromatin organization [8–11]. NL is composed of A and B type lamins along with inner nuclear membrane (INM) proteins [12], and together with mediator proteins such as Barrier-to-Autointegration Factor (BAF) and Heterochromatin Protein 1 (HP1), facilitate attachment of chromatin to NL [13,14]. Additionally, these interactions have been proposed to form specific chromatin organization that opposes transcriptional activity [15]. The association between LMNB receptor and LMNA/C mediates peripheral heterochromatin attachment in a wide variety of mammalian cells [16]. Any perturbation in such organization leads to a complete loss of peripheral heterochromatin and developmental abnormalities [17].

Recent studies demonstrated that the Lamina associated domains (LADs) enriched in H3K9 methyl (me2/me3) marks contact NL via association with LMNB1 [7,18–20]. These interactions are highly stochastic in nature and are dependent on H3K9me2 activity governed by G9a/EHMT2. Accordingly, G9a/EHMT2 promotes LAD formation and its loss leads to the opposite effect [18]. Similar to humans, H3K9 methylation is important for heterochromatin positioning in *C. elegans* [21], as depletion of H3K9 methyltransferases Met2 and Set-25 (mammalian SETDB1 and G9a /EHMT2 homolog) leads to detachment of large gene-array from peripheral heterochromatin. Altogether, loss of lamins and INM proteins or H3K9me2 activity leads to peripheral heterochromatin defects, however, whether there is a link between these common consequences remains unknown. Here we establish EHMT proteins as a common module that governs heterochromatin tethering via histone dependent (H3K9me2) and independent mechanisms (by directly regulating LMNB1 methylation).

## Results

### EHMTs associates with LMNB1

To identify the novel non-histone interactors of EHMT proteins, endogenous EHMT1 was immunoprecipitated (IP) and unique bands in the EHMT1 pull down were subjected to LC/MS analysis. We found LMNB1 and histone proteins as interactors of EHMT1 (Fig 1A). Mass spectrometry data was validated by sequential IP reactions of endogenous EHMT1 and LMNB1 proteins (Fig 1B). This interaction was also found using nuclear extracts from human dermal fibroblasts (HDFs) (Fetal derived unless mentioned otherwise), suggesting that it is not cell type specific interaction (Fig Supplementary Fig 1A). Consistent with previously published reports, we also detected HP1 in association with EHMT and LMNB1 (Supplementary Fig 1A). The absence of Ash2l (a member of H3K4 methyltransferase complex) in the IP-EHMT1 or IP-LMNB1 confirmed the specificity of IP reaction (Supplementary Fig 1A). Mapping experiments (*in vivo* and *in vitro*) to identify the LMNB1 interacting domain of EHMT1, revealed that the SET domain of EHMT1 is sufficient to bind to LMNB1 (Supplementary Fig 1B and C; Fig 1C). These results confirmed that the EHMT1/2 directly associates with LMNB1 via its SET domain.

**Figure 1.**
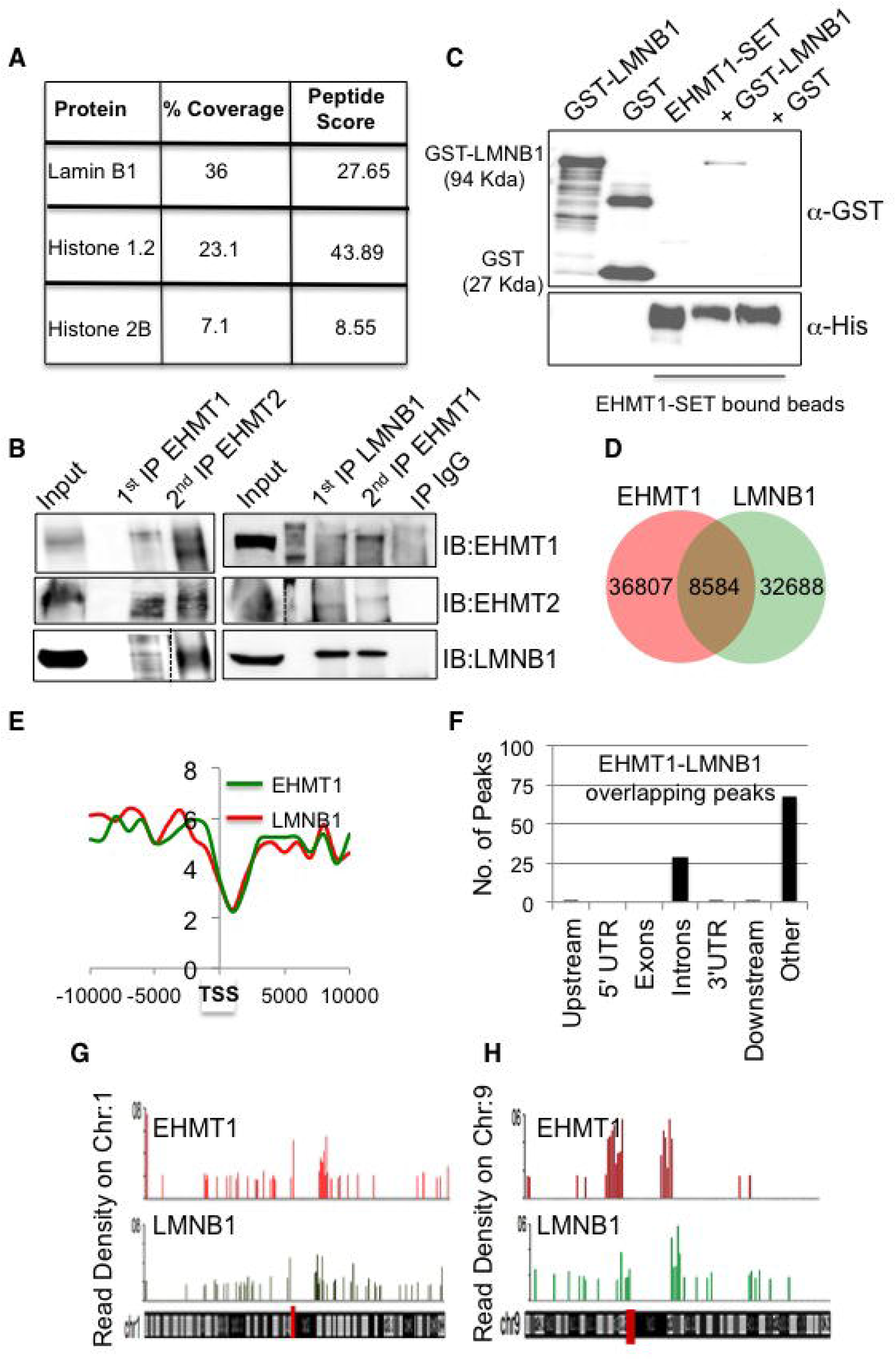
EHMT1, EHMT2, and LMNB1 are members of the same complex. **A** EHMT1 interacting proteins identified by mass spectrometric analysis with details indicating coverage and peptide score. **B** Sequential IP in HEK293 cells demonstrating EHMT1, EHMT2, and LMNB1 are a part of the same complex. **C** LMNB1 interacts with EHMT1 via SET domain. Recombinant GST or GST-LMNB1 was incubated with Ni-NTA bound His-EHMT1 SET protein. Post washing eluents were loaded for immunoblotting using GST or His antibody. Recombinant pure proteins GST-LMNB1 (lane 1), GST (lane 2), EHMT1-SET (lane 3) were used as controls. **D** Venn diagram showing unique and overlapping reads obtained from EHMT1 and LMNB1 ChIP-Sequencing. **E** Composite profile of EHMT1 and LMNB1 read density around the transcription start site (TSS). **F** Genomic distribution of EHMT1 and LMNB1 peaks. The majority of binding sites obtained were enriched in an intronic region or distal regions from a gene. **G, H** Representative figure showing normalized ChIP-seq read density (above 1.5-fold over expected) of EHMT1 and LMNB1 in 1MB bin for chromosome 1 & 9.

Both EHMT and LMNB1 are known to interact with chromatin independently or via mediator proteins [17,22,23]. To identify EHMT1-LMNB1 co-bound regions in the genome, we performed ChIP-Seq analysis. Individually, EHMT1 and LMNB1 occupied 36807 and 32688 number of peaks respectively and 8584 peaks were co-bound by EHMT1 and LMNB1 (Fig 1D). A majority of EHMT1 and LMNB1 reads were distributed on non-TSS regions such as introns and gene-poor regions (represented as “others”) (Fig 1E and F), whereas only 1.5% of reads were found on the upstream region of genes (Fig 1F). Functional category analysis of the genes occupied by EHMT1, LMNB1 or EHMT1-LMNB1 revealed enrichment of genes regulating transcription, signal transduction and cell adhesion (Supplementary Fig 1D-F).

H3K9me2 deposited by EHMT1/2 complex demarcates heterochromatin, particularly non-genic regions and is prevalent in “other regions” such as gene deserts, pericentromeric and subtelomeric regions, with little being observed at individual active or silent genes. Detailed analysis of read densities performed on individual chromosomes identified a significant correlation (p<0.001) between EHMT1 and LMNB1 localization. Interestingly, we observed higher ChIP-seq read density onto subtelomeric, and around centromeric regions (indicated as a red line on the chromosome) that are preferentially maintained in the silent state (Fig 1G, H, and Supplementary Fig 1G). These data suggested that EHMT1-LMNB1 associates on gene-poor areas that are the critical determinants of higher-order chromosome structure at the NP.

### EHMTs methylate LMNB1

Next, we tested if LMNB1 is a substrate for methylation by the EHMT enzymes. Using an *in vitro* fluorometric methyltransferase assay we demonstrate an increase in fluorescence upon incubation of EHMT1-SET domain with GST-LMNB1 in presence of S-adenosyl methionine (SAM) (Supplementary Fig 2A). To confirm that EHMT proteins indeed methylate LMNB1, we used lysine methyl-specific (Methyl-K) antibody to probe for methylated LMNB1. Purified LMNB1 C terminus protein containing the rod domain and tail domains (LMNB1-CT) (Supplementary Fig 2B) was used in this assay. Towards this, we performed *in vitro* methyltransferase assay using different concentrations of LMNB1 and incubated with an equimolar ratio of the EHMT1/2-SET domain in presence or absence of SAM. When products of these reactions were immunoblotted using the Methyl-K antibody, specific methylation signal was observed upon incubation of LMNB1 with EHMT1/2-SET in presence of SAM (Fig 2A and B).

**Figure 2.**
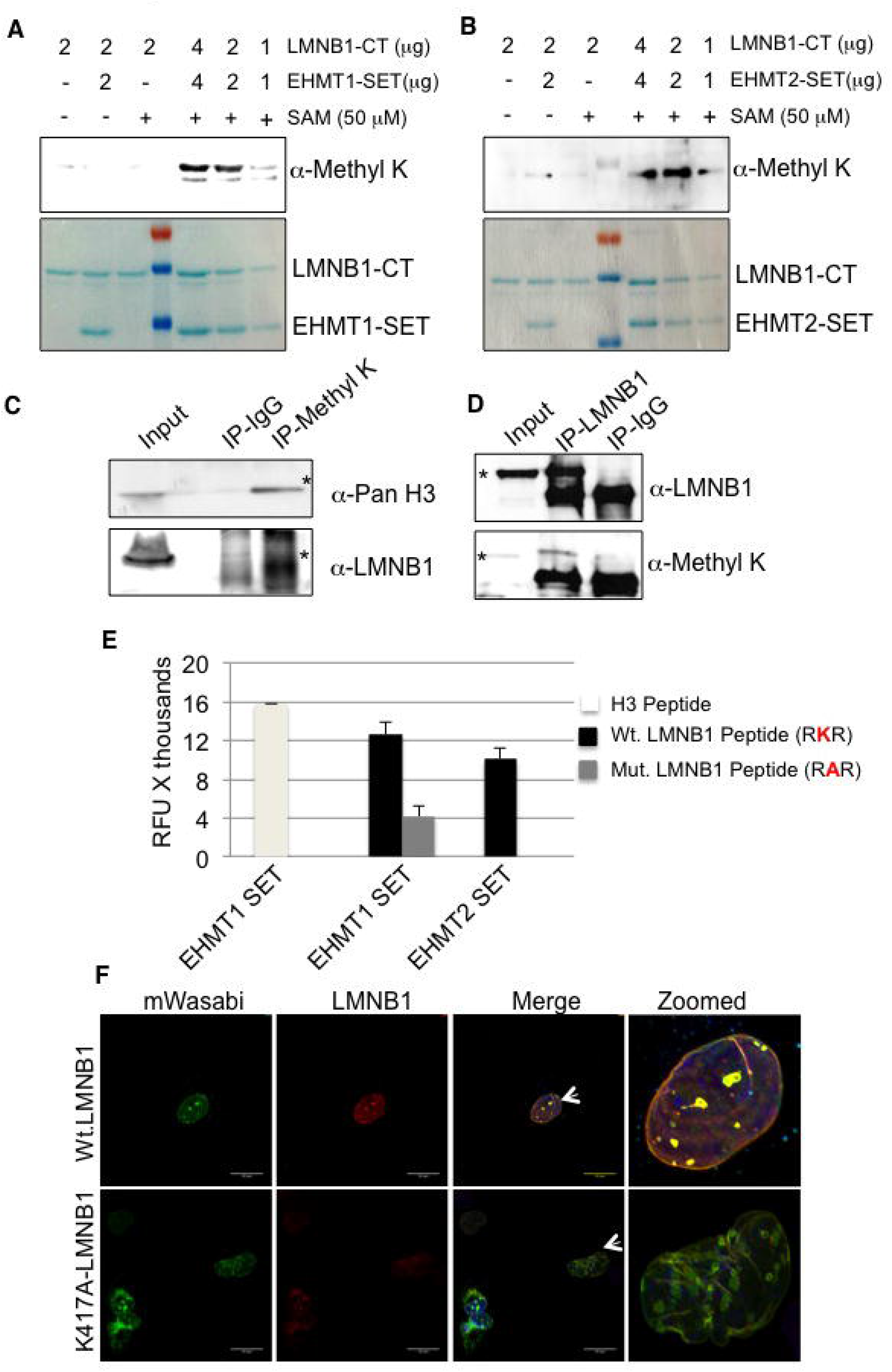
EHMT1 and EHMT2 methylate LMNB1 at C-terminus. **A, B** Western blot probed with the anti-Methyl-K antibody. Lanes to the left of the ladder are control reactions containing 2 μg LMNB1-CT + 50 μM SAM (lane 1) or 2 μg of EHMT1-SET/EHMT2-SET + 2 μg LMNB1-CT (lane 2) and 2 μg LMNB1-CT only (lane 3). Lanes to the right of the ladder are methyltransferase reactions containing 1, 2 and 4 μg of LMNB1-CT and EHMT1-SET/EHMT2-SET along with 50 μM SAM. Lower panel: Coomassie stained gel representing all the reactions mentioned above. **C, D** Nuclear lysates from HEK-293 were immunoprecipitated with anti-Methyl-K or anti-LMNB1 antibody followed by western blotting with anti-Pan H3, anti-LMNB1 and anti-Methyl-K antibodies. Asterisk indicates the band of interest. **E** Reduced methylation of LMNB1 by EHMT1 and EHMT2 upon single amino acid substitution from lysine (RKR) to alanine (RAR). Mutant LMNB1 peptides were synthesized and subjected to methyltransferase reaction containing EHMT1-SET and EHMT2-SET with SAM as a methyl donor. H3 peptide was used as a positive control for the reaction. **F** Immunostaining for LMNB1 in fetal HDFs transfected with Wt.LMNB1 and K417A-LMNB1 constructs. (Scale bar: 20μm). Arrows indicate the cells zoomed in the far-right image presented.

Further, to examine if LMNB1 is methylated *in vivo,* we performed anti-Methyl-K or anti-LMNB1 IPs from HEK nuclear lysates. Products of IPs were split into two halves and probed with either anti-LMNB1 or anti-Methyl-K specific antibodies. Several lysine residues are methylated on the H3 tail, detecting the histone signal in IP-Methyl-K confirmed the specificity of IP reaction and served as a positive control (Fig 2C). The presence of a LMNB1 band in the same Methyl-K IP samples indicated the presence of endogenous methylated LMNB1 (Fig 2C). In reciprocal IP reaction in which LMNB1 was IP’ed and probed with Methyl-K antibody, identification of Methyl-K signal in the IP-LMNB1 confirmed LMNB1 methylation *in vivo* (Fig 2D). It has been reported that EHMT2 is capable of methylating lysine on dipeptide Arg-Lys (RK) sequence of non-histone proteins [24]. We synthesized peptides for such motifs present at the C-terminus of LMNB1 and identified K417 as the methylation site targeted by EHMT1 and EHMT2 (Fig 2E). K417A peptide mutation abolished methylation of LMNB1 (Fig 2E). To investigate the function of methylated LMNB1 *in vivo,* we mutated the 417K residue to alanine (K417A) in the wild-type (Wt.) LMNB1 construct. As opposed to Wt.LMNB1, which was localized at the NP, much of K417A-LMNB1 was accumulated in the nucleoplasm (Fig 2F and Supplementary Fig 2C-E). We also observed aggregates of mutant LMNB1 transported into the cytoplasm, and was accompanied by abnormal nuclear morphology (Fig 2F and Supplementary Fig 2C, 2F). Co-staining with LMNB1 antibody showed localization of endogenous LMNB1 and overexpressed Wt.LMNB1 at the NP in Wt.LMNB1 expressing cells (Fig 2F). However, in mutant-LMNB1 expressing HDFs, endogenous LMNB1 was localized in K417A-LMNB1 aggregates indicating a dominant negative function of the mutant protein (Fig 2F). Further mislocalization of LMNA/C in the aggregates of mutant-LMNB1 (Supplementary Fig 2G) indicated LMNB1 methylation is critical for maintaining the NL meshwork composition at the periphery. Based on these data we speculated that the methylation modification prevents the degradation of LMNB1 and confers protein stability.

### EHMTs regulate LMNB1 levels

To test the consequence of the loss of EHMTs on LMNB1 levels we used shRNAs and achieved approximately 60% and 70% depletion of EHMT1 and EHMT2, respectively (Supplementary Fig 3A-C). Reduced expression of EHMT2 in shEHMT1 cells and vice-versa indicated reciprocal stabilization of these proteins within the heteromeric complex (Supplementary Fig 3A-C). Further reduced levels of LMNB1 protein in shEHMT1 and shEHMT2 cells (Fig 3A) confirm the regulation of LMNB1 by EHMTs. Moreover, shEHMT1 and shEHMT2 cells exhibited a significant distortion of nuclear morphology (Fig 3B, C, and Supplementary Fig 3D). Recent studies demonstrate that the B-type lamins are long-lived proteins [25]. Therefore, loss of such proteins is a combination of transcriptional inhibition and protein degradation. Thus, we tested if EHMT proteins regulated LMNB1 expression transcriptionally. Our data demonstrate approximately 60% loss of LMNB1 transcript upon depletion of EHMT proteins (Fig 3D). Overall our results demonstrate that EHMT1 and 2 regulate LMNB1 expression transcriptionally and directly via post-translational modification.

**Figure 3.**
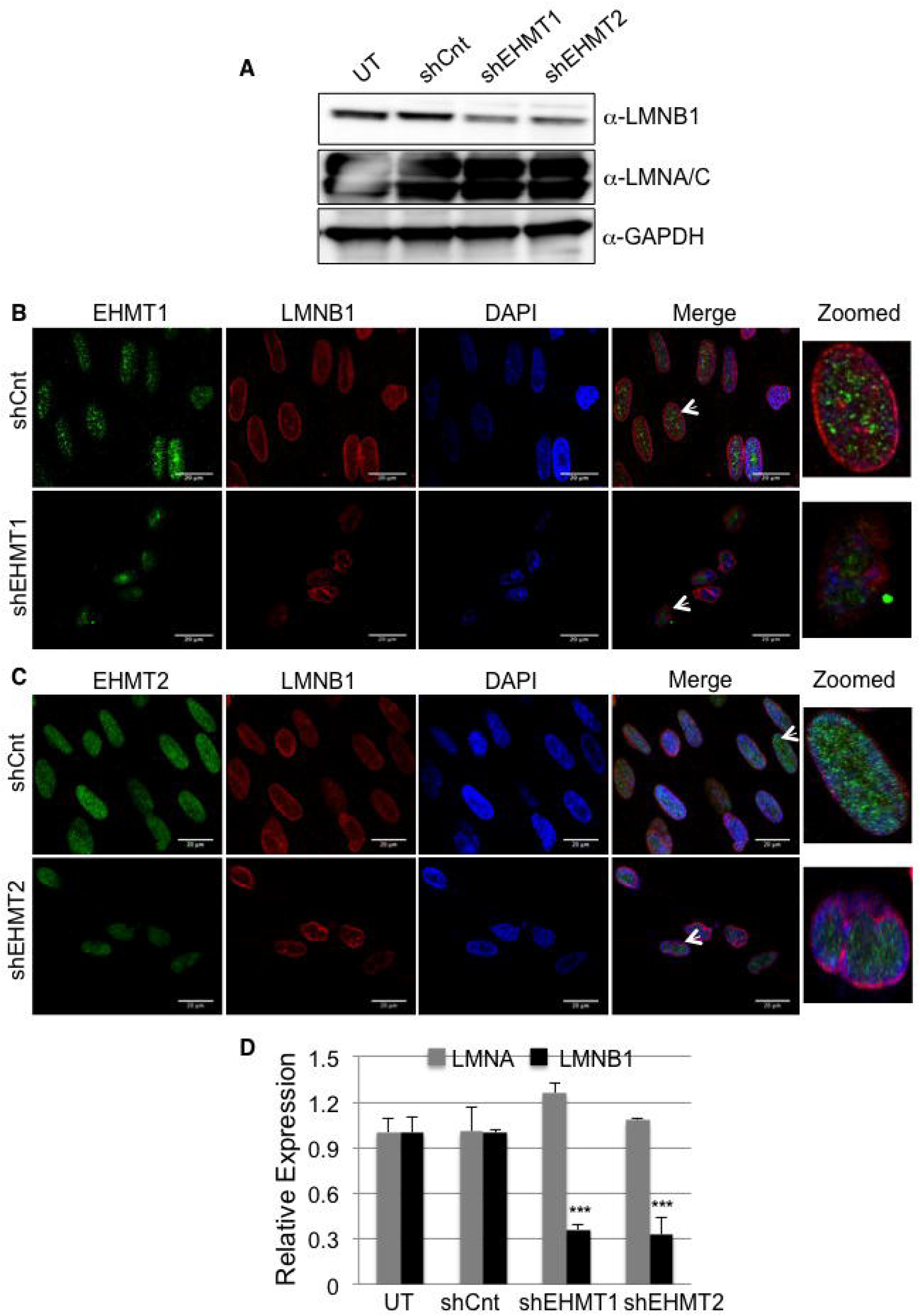
EHMT1 and EHMT2 knockdown results in reduced LMNB1 levels. **A** Western blot analysis for LMNB1 and LMNA/C in fetal HDFs transduced with shEHMT1 and shEHMT2 virus. Untransduced (UT) and control shRNA (shCnt) were used as controls. **B, C** Fetal HDFs were transduced with shCnt, shEHMT1 and shEHMT2 virus. Post transduction cells were immunostained for EHMT1 or EHMT2 and co-stained for NL using LMNB1 antibody. Distortion of nuclear architecture was seen upon loss of EHMT1 and EHMT2 proteins. (Scale bar: 20μm). Arrows indicate the cells zoomed in the far-right image presented. **D** Relative expression of LMNA and LMNB1 in fetal HDFs upon EHMT1 and EHMT2 knockdown compared to UT/shCnt cells. (n=3) For LMNB1: UT vs shEHMT1, ***p<0.0001 and UT vs shEHMT2, ***p<0.0001. (One Way ANOVA, post-hoc test: Bonferroni’s multiple comparison test).

### EHMT mediated histone and non-histone methylation influences peripheral heterochromatin organization

H3K9 methylation and LMNB1 are the critical determinants for the formation of the LADs at the NP [7,18–20]. To test the requirement of EHMT mediated H3K9 dimethylation in tethering peripheral heterochromatin, we depleted EHMTs and monitored the co-localization of H3K9me2 with LMNB1. Global H3K9me2 was decreased by 50% in shEHMT1 and 80% in shEHMT2 cells (Supplementary Fig 4A and B), confirming that EHMT2 is the predominant HMTase among EHMT proteins. Though H3K9me2 is primarily localized to the NP in shCnt and shEHMT1 HDFs, there was a significant loss of enrichment of H3K9me2 in shEHMT2 cells (Supplementary Fig 4C-E and Fig 4A).

**Figure 4.**
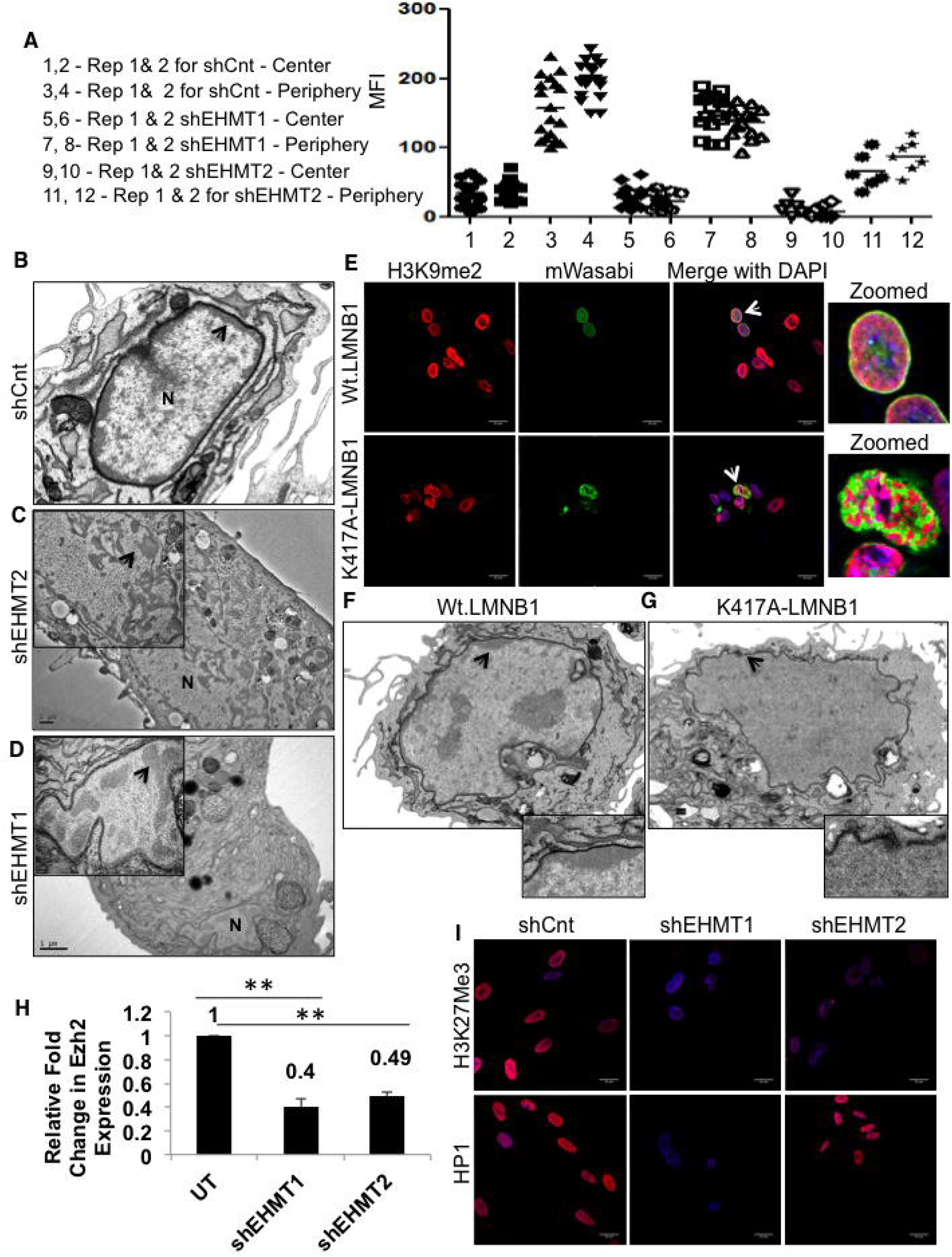
Reduction in EHMT1 and EHMT2 results in detachment of peripheral heterochromatin. **A** MFI for H3K9me2 staining in fetal HDFs transduced with shEHMT1 and shEHMT2. MFI has been represented as center vs. periphery of the nuclei and compared with respect to shCnt cells. Each biological replicate is represented as spread of technical replicates around the mean value. **B, C, D** Fetal HDFs were transduced with shCnt, shEHMT1 and shEHMT2 virus. TEM was performed to visualize the heterochromatin in EHMT1 and EHMT2 knockdown cells. Peripheral heterochromatin was intact at the NP beneath the NL in control cells while this distribution was drastically altered upon knockdown of EHMT1 and EHMT2. (Scale bar: 1μm). Arrows indicate the heterochromatin regions. **E** Immunostaining for H3K9me2 in fetal HDFs expressing Wt.LMNB1 or K417A-LMNB1 mutant plasmids. (Scale bar: 20μm). Arrows indicate the cells zoomed in the far-right image presented. **F, G** Fetal HDFs transfected with Wt.LMNB1 and K417A-LMNB1 plasmids were processed for electron microscopy. Cells transfected with Wt.LMNB1 plasmid showed intact peripheral heterochromatin while K417A-LMNB1 transfected cells showed loss of heterochromatin with nuclear envelope breaks. (Scale bar: 1μm). Arrows indicate the zoomed area presented in the inset below. **H** qRT-PCR to validate expression of EZH2 in UT, shEHMT1 and shEHMT2 transduced HDFs. (n=3), UT vs. shEHMT1 (**p=0.0067), UT vs. shEHMT2 (**p=0.0021). (One sample t-test, two-tailed). **I** Immunostaining for H3K27me3 mark and HP1 protein in fetal HDFs transduced with shCnt, shEHMT1, and shEHMT2. Knockdown of EHMT1 and EHMT2 leads to a reduction in the H3K27me3 mark, which corroborates with decreased Ezh2 expression while HP1 expression was reduced only in shEHMT1 cells. (Scale bar: 20μm)

Next, we performed EM to investigate the status of heterochromatin in EHMT depleted cells. shCnt transduced HDFs exhibited a layer of electron dense peripheral heterochromatin just beneath the nuclear envelope (NE) (Fig 4B). Knockdown of EHMT2 led to the partial disruption of heterochromatin from the periphery to the interior of the nucleus (Fig 4C). This result was correlated with redistribution of H3K9me2 marks towards the interior of the nucleus. In shEHMT1 HDFs, we noticed near complete detachment of peripheral heterochromatin and a distorted NE (Fig 4D). We also detected floating islands of heterochromatin in the nuclei. Interestingly, the severity of heterochromatin detachment and compromised NE integrity were unique to shEHMT1 knockdown wherein H3K9me2 activity was modestly affected.

We also looked at the effects of pharmacological inhibition of H3K9me2 on overall heterochromatin positioning and nuclear distortion. HDFs treated with BIX 01294 showed 40% less H3K9me2 staining compared to controls (Supplementary Fig 4F-H) with no changes in the LMNB1 methylation levels (Supplementary Fig 4H). Unlike EHMT depleted cells, we did not notice any significant changes in the nuclear morphology of BIX treated cells (Supplementary Fig 4F and I). EM imaging indicated modest changes in heterochromatin anchorage (Supplementary Fig 4I). Interestingly addition of BIX did not influence EHMT mediated methylation of LMNB1 in fluorometric methyltransferase assay using recombinant LMNB1 protein (Supplementary Fig 4J).

To obtain a clearer picture of the role of LMNB1 methylation, HDFs were transfected with either the Wt.LMNB1 or K417A-LMNB1 construct. Overexpression of Wt. or mutant LMNB1 did not alter the levels of H3K9 dimethylation (Supplementary Fig 4K-L). Immunostaining for H3K9me2 revealed the colocalization of H3K9me2 with LMNB1 at the NP in Wt.LMNB1 overexpressing cells (Fig 4E upper panel). In mutant-LMNB1 expressing cells, the peripheral distribution was severely compromised and we noticed segregation of LMNB1 and H3K9me2 aggregates in the nucleoplasm (Fig 4E lower panel and Supplementary Fig 4M). EM analysis revealed that the nuclei of mutant-LMNB1 expressing cells were devoid of peripheral heterochromatin and displayed a ruptured nuclear envelope (Fig 4F-G). This data suggests that the mislocalization of K417A LMNB1 could be responsible for the loss of peripheral distribution of chromatin. Nonetheless, our results demonstrate the significance of EHMT mediated LMNB1 methylation in stabilizing LMNB1 at the NP and heterochromatin organization.

To investigate the additional changes that could influence heterochromatin organization in EHMT depleted cells we profiled gene expression changes in shEHMT1 and shEHMT2 HDFs. RNA-Seq analysis identified overlapping and unique genes that were regulated by EHMT1 or EHMT2 (Supplementary Fig 5A, Supplementary Table 1 and 2), which were enriched for genes involved in the cell-cycle, homeostasis and axon guidance etc. (Supplementary Table 3). Interestingly, significantly high numbers of pathways were distinctly regulated by EHMT1 and EHMT2 (Supplementary Table 3). The varying degree of peripheral heterochromatin detachment in shEHMT1 vs. shEHMT2 HDFs led us to investigate the number of chromatin modifiers that were altered upon EHMT depletion. While there were overlapping chromatin modifiers that changed in response to EHMT depletion, EHMT1 loss reduced expression of repressive PRC1 components (Supplementary Table 4). On the contrary EHMT2 reduction predominantly influenced the expression of KDMs, SIRT6 and SETD1 proteins (Supplementary Fig 5B, Supplementary Table 4). Functional validation of the loss of PRC members was performed by assaying EZH2 transcript and H3K27 methyl marks. Reduced EZH2 expression and H3K27me3 staining in shEHMT1 and shEHMT2, indicates that knockdown of EHMTs affects PRC protein expression (Fig 4H and I upper panel). Interestingly, we detected the loss of HP1 only in shEHMT1 cells but not in shEHMT2 HDFs (Fig 4I, lower panel). Overall our results revealed the commonalities and differences upon individual knockdown between two structurally similar proteins that contribute to the phenotype of peripheral heterochromatin detachment.

### Sequential loss of EHMT proteins correlates with diminished peripheral heterochromatin organization during physiological aging

The cellular phenotypes induced by EHMT1 knockdown impinge on known molecular hallmarks of cellular aging [26]. These observations prompted us to survey if age-related genes were affected in response to EHMT depletion. Comparison of RNA-Seq profile obtained from shEHMT1 and shEHMT2 HDFs identified approximately 30% of aging specific genes (Total 108 of 307 genes listed in the aging database) were altered in response to depletion of EHMT (Supplementary Fig 6A and Supplementary Table 5). PCR analysis verified that genes linked to senescence, aging and diabetes are differentially regulated upon EHMT depletion (Supplementary Fig 6B and C).

To test if EHMTs and H3K9me2 are indeed involved in regulating the aging process, we monitored the expression of EHMT1 and EHMT2 in HDFs derived from individuals of different age groups. As previously reported histone methyl marks H3K27me3, H3K9me3 and HP1 are diminished in aged HDFs (Supplementary Fig 6D and E). Further investigation into levels of EHMT proteins revealed a decline of EHMT1 and EHMT2 in an age-dependent manner (Fig 5A-D). Compared to fetal cells, 18Y old HDFs exhibited decline in EHMT2 and preceded the loss of EHMT1 (Fig 5B and C). An investigation of the status of H3K9me2 methylation demonstrated its decline in aging fibroblasts (Fig 5E and F). An equally important observation was that the preferential localization of H3K9me2 was noticed at the NP in fetal HDFs, while these marks were distributed in the nucleoplasm in 18Y and 65Y HDFs (Fig 5F and Supplementary Fig 7A-D). This data correlates with our observations that reduction of EHMT2 in fetal-HDFs results in the re-localization of H3K9me2 methyl marks to the nucleoplasm (Fig 4A and Supplementary Fig 4C).

**Figure 5.**
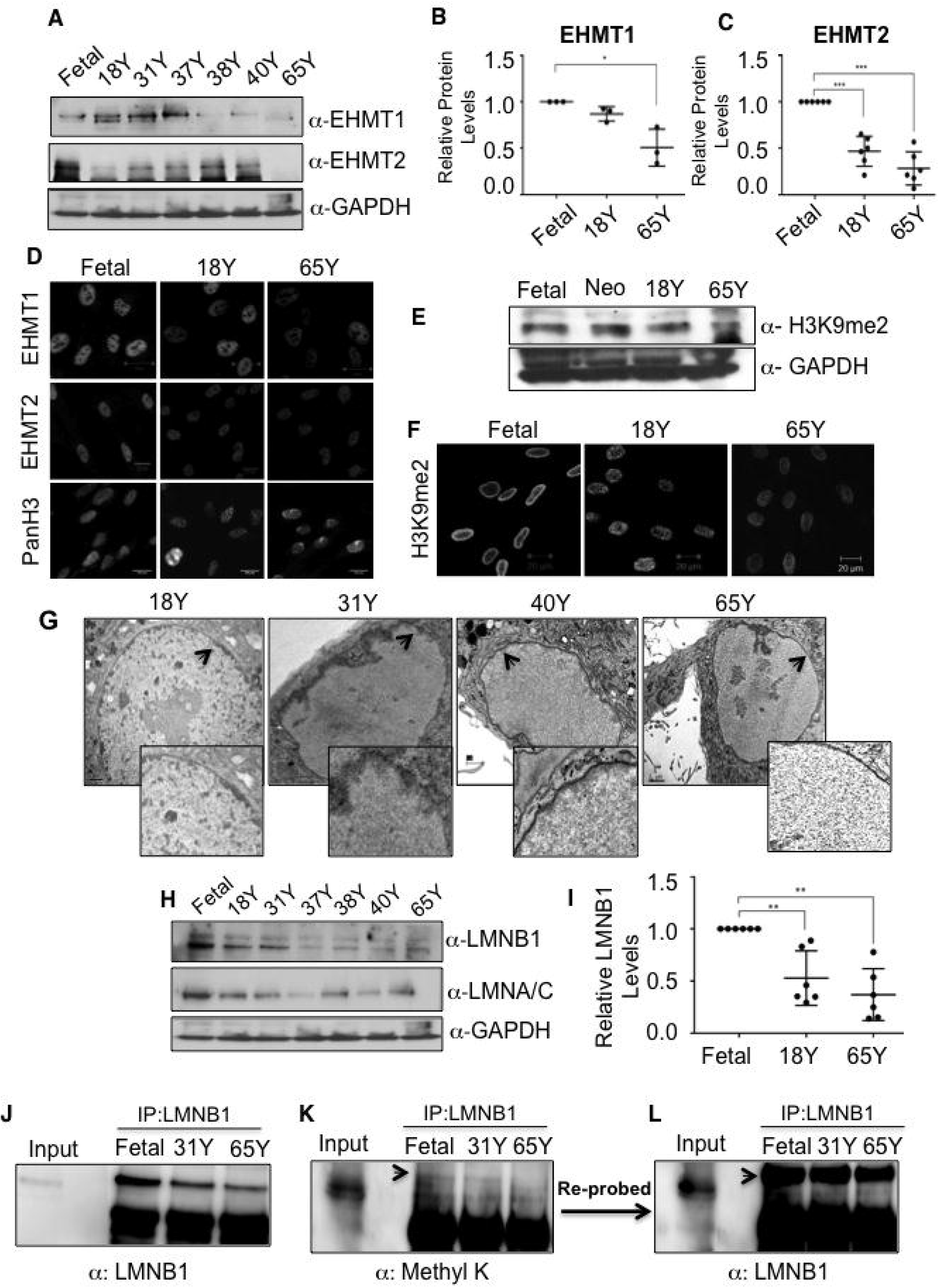
Depletion of EHMT1, EHMT2 and LMNB1 in an age dependent manner. **A** Western blot analysis for EHMT1, EHMT2 and GAPDH in HDF cell lysates derived from various age groups. **B**, Quantification of EHMT1 protein in HDFs of indicated age groups. One Sample t test (n=3). For fetal vs 65 (*p<0.05), Fetal vs 18Y and Fetal vs 31Y are not significant. **C**, Quantification of EHMT2 protein expression in HDFs of indicated age groups. One Sample t test (n=6). For Fetal vs 18Y (***p=0.0004), and Fetal vs 65Y (***p=0.0002). **D** Immunostaining for EHMT1, EHMT2 and Pan H3 in fibroblasts from indicated age groups (Scale bar: 20μm). **E, F** Western blot and immunostaining (Scale bar: 20μm) analysis for H3K9me2 in indicated age groups. **G** Representative TEM images for nuclei of 18Y, 31Y, 40Y and 65Y HDFs. Arrows indicate peripheral heterochromatin. Inset is the zoomed version of the same images. (Scale bar: 1μm). **H** Western blot analysis for LMNA/C and LMNB1 and in various age groups HDFs. GAPDH was used as an internal control. **I** Quantification of LMNB1 protein expression in HDFs of indicated age groups. One sample t test (n=6). For Fetal vs 18Y (**p=0.0069) and Fetal vs 65Y (**p=0.0016). **J,K,L** Reduced levels of methylated LMNB1 during physiological aging. Nuclear extracts from indicated age groups were subjected to IP using LMNB1 antibody. IPed material was divided into halves, Blot 1 (left,J) was probed with anti LMNB1 antibody, blot 2 was probed with anti-Methyl-K (middle, K) antibody. Antimethly-K blot was reprobed with LMNB1 antibody to demonstrate the methyl band indicated is indeed LMNB1 (Right, L). 80 μg of fetal HDFs derived nuclear lysate was used as input control.

We next examined whether progressive loss of EHMTs and altered distribution of H3K9me2 impacts heterochromatin organization in aging cells. Compared to fetal cells, in 18Y and 31Y HDFs, heterochromatin was still retained at the periphery where there was a substantial loss of EHMT2 (Fig 5G and A). We also observed a redistribution of peripheral heterochromatin, which correlated with redistribution of H3K9me2 marks (Fig 5G and Fig 4B). Further, there was a gradual loss of peripheral heterochromatin in 40Y old cells with complete depletion observed in 65Y aged nuclei (Fig 5G). Interestingly complete loss of peripheral heterochromatin organization was correlated with loss of EHMT1 and EHMT2 in aged cells as shown in Fig 5A-C.

Since both EHMTs were downregulated during aging we explored the regulation of EHMT1 and EHMT2 transcripts and proteins. The EHMT1 transcript was downregulated by 10% from fetal to 18Y HDFs and was further downregulated by 90% from 18Y to 65Y stage (Supplementary Fig 7E). This data clearly indicated that EHMT1 expression is controlled transcriptionally during aging. On the contrary, only 30% of EHMT2 was regulated transcriptionally (20% downregulation from fetal to the adult state, with a further decline by 30% from adult to aged cells) (Supplementary Fig 7E). Since a small amount of EHMT2 was regulated transcriptionally, there seemed a strong possibility of post-translation regulation during aging. Hence, to investigate if EHMT2 protein levels are regulated via ubiquitin-proteasome pathway (UPS), we treated 18Y and 65Y old HDFs with the proteasome inhibitor MG-132. In 18Y old cells, EHMT2 protein levels were increased to 2-fold upon MG-132 treatment, whereas no such increase was noticed in EHMT1 levels (Supplementary Fig 7F and G). Interestingly, EHMT2 levels could not be rescued in 65Y HDFs (Supplementary Fig 7F), indicating that the blockade of UPS activity can restore the EHMT2 degradation only in early age group and such mechanisms do not operate in aged cells wherein EHMT2 is already drastically low. Taken together our data on the loss of EHMT1 either as a consequence of physiological aging or by forced depletion in fetal HDFs implicates EHMTs in heterochromatin organization at the NP.

Low levels of LMNB1 have been observed in senescent cells and fibroblasts derived from Progeria patients [27]. In this study, expression analysis of nuclear lamins in intrinsically aged cells showed decline in LMNA/C from fetal to 18Y age and then the protein levels remained constant in further age groups (Fig 5H). Contrary there was a dramatic reduction of LMNB1 starting in the 18Y age group with a significant loss at 65Y (Fig 5H and I). Diminishing levels of LMNB1 were correlated with the reduction in EHMT2 protein with drastic loss upon depletion of EHMT1 protein in 65Y cells (Fig 5A-D). This is consistent with the data in Figure 3 where we found that EHMT proteins directly regulate levels of the LMNB1 protein.

Next, we investigated whether diminishing perinuclear heterochromatin organization in aging nuclei is a result of the loss of EHMT1, EHMT2 and LMNB1 interaction. Towards this, we performed IP experiments using Fetal, 18Y and 65Y HDF nuclear extracts. Our results revealed an association between EHMT2 and LMNB1 occurred in fetal cells and was highly diminished in adult and aged cells (Supplementary Fig 2H). On the contrary, EHMT1 associated with LMNB1 in all the age groups and the interaction was reduced gradually in age-dependent manner (Supplementary Fig 7I). The complete absence of perinuclear heterochromatin in 65Y-aged nuclei corresponded to over 80% reduction in the interaction between EHMT1 and LMNB1. These data indicated that EHMT1 and LMNB1 association is involved in maintaining peripheral heterochromatin in aging fibroblasts.

Further, we tested the status of LMNB1 methylation during the physiological aging and found the reduced intensity of a methylated LMNB1 signal from Fetal to 31Y with virtually no methylation in 65Y-aged cells (Fig 5J-L). Taken together our results indicate that the loss of peripheral heterochromatin in EHMT depleted cells or aged cells occur due to loss of H3K9 activity coupled with the loss of LMNB1, which are critical determinants of peripheral heterochromatin anchorage.

### Overexpression of EHMT proteins rescues peripheral heterochromatin defect in aged cells

To test if depletion of EHMT proteins is indeed responsible for the loss of peripheral heterochromatin in aged cells we transfected full-length V5 tagged EHMT1 or Flag-EHMT2 (set domain) plasmids in 65Y HDFs. Immunostaining using V5 or Flag antibodies confirmed the overexpression of EHMT1 and EHMT2 proteins (Fig 6A). EHMT1 levels enhanced the expression of LMNB1 but the Flag-SET of EHMT2 had no effect on LMNB1 levels (Fig 6A middle and lower panel and Supplementary Fig 8A and B).

**Figure 6.**
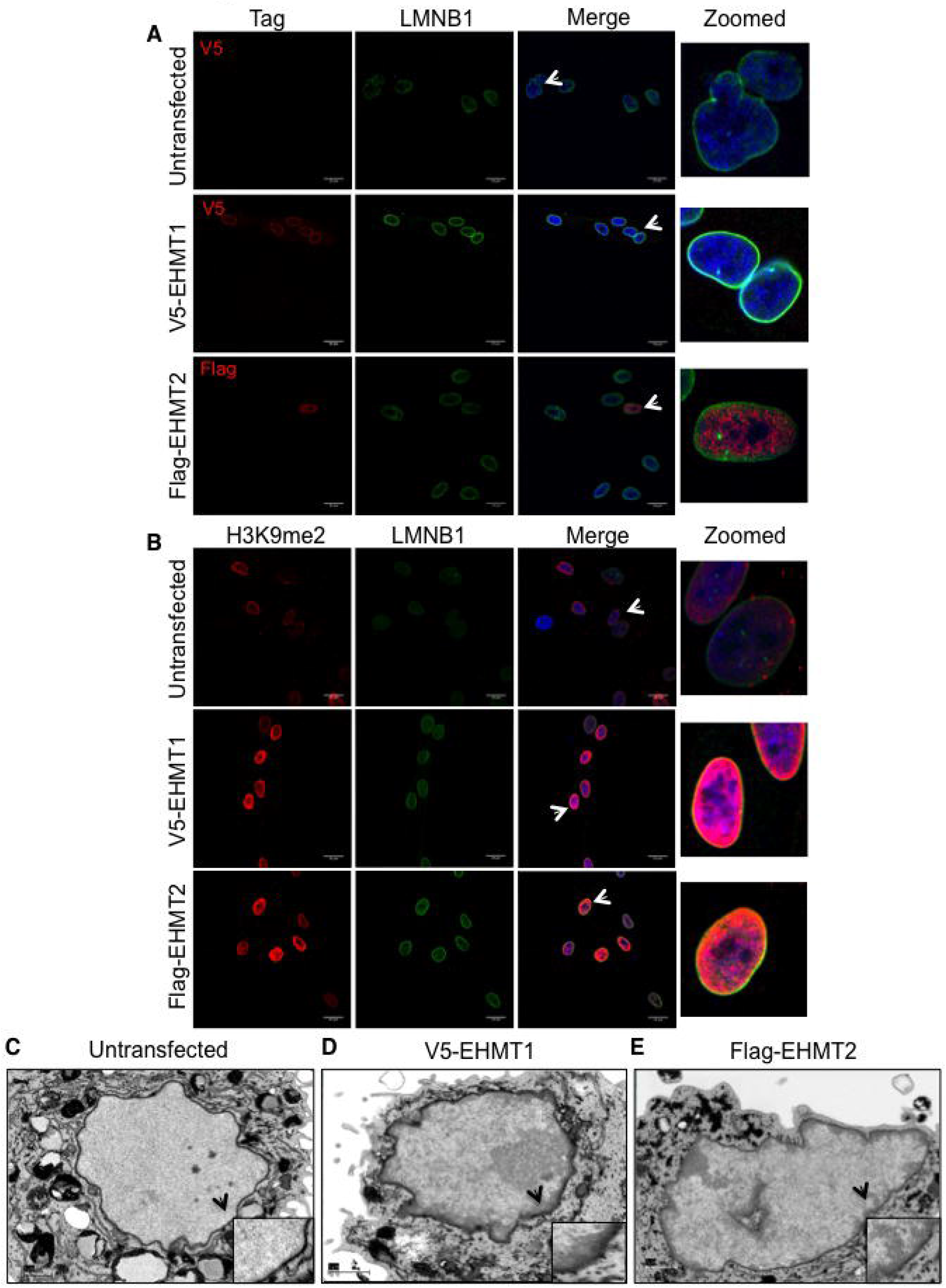
Overexpression of EHMT1 and EHMT2 restores peripheral heterochromatin organization in aged cells. **A, B** Immunostaining using V5 or Flag antibodies along with LMNB1 in 65Y HDFs transfected with V5-EHMT1 and Flag-EHMT2 overexpression constructs. Immunostaining using H3K9me2 antibody along with LMNB1 in 65Y HDFs transfected with V5-EHMT1 and Flag-EHMT2 overexpression constructs. (Scale bar: 20μm) Arrows indicate the cells zoomed in the far-right image presented. **C, D, E** 65Y Old HDFs overexpressing EHMT1 and EHMT2 were processed for electron microscopy. Overexpression of EHMT1 and EHMT2 causes restoration of peripheral heterochromatin in old cells compared with control cells (Scale bar: 1μm). Arrows indicate the area zoomed and presented in the inset format.

We further examined H3K9me2 localization and organization of heterochromatin in EHMT1 and EHMT2 overexpressing cells. In both the cases H3K9me2 was co-localized with LMNB1 at the NP (Fig 6B). Consistent with this, EM imaging revealed peripherally organized heterochromatin upon EHMT1 and EHMT2 overexpression, which was completely absent in untransfected aged cells (Fig 6C-E). These results demonstrate that the reduction of EHMT proteins contributes to the loss of peripheral heterochromatin organization during aging and this defect can be reversed upon re-expression of EHMT proteins.

To investigate the contribution of LMNB1 methylation in reversing peripheral heterochromatin tethering in aged cells we co-expressed wild-type or mutant-LMNB1 with V5-EHMT1 or Flag-EHMT2. As expected Wt.LMNB1 localized with H3K9me2 at the NP in EHMT1 overexpressing cells (Supplementary Fig 8C, upper panel). Instead, H3K9me2 showed aggregated staining in the nucleoplasm and did not localize with mutant-LMNB1 (Supplementary Fig 8D and E) thereby implicating a role for methylated LMNB1 in organizing heterochromatin to the NP.

### Proliferation rate of HDFs correlate with EHMT expression

Previous reports have demonstrated that the reduction in LMNB1 expression results in reduced proliferation and induction of premature senescence [28,29]. Therefore, we tested if the reduction of LMNB1 expression upon loss of EHMTs leads to altered proliferation. Interestingly, shEHMT1 & 2 showed a significant reduction in cell number (Fig 7A and B). Cell-cycle analysis revealed a small fraction of EHMT2 transduced cells (8.4%), were in SubG1 phase compared to shCnt and shEHMT1 HDFs (Supplementary Fig 9A-C and Fig 7C), indicating a small but significant apoptosis in shEHMT2 cultures. This observation is supported by transcriptome data showing the upregulation of apoptotic genes upon knockdown of EHMT2 (Supplementary Table 2 and 3). shRNA mediated loss of EHMT proteins also induced senescence program. We observed higher number (approximately 15 fold) of senescent cells in shEHMT1 in comparison to shEHMT2 (2 fold) (Fig 7D and E).

**Figure 7.**
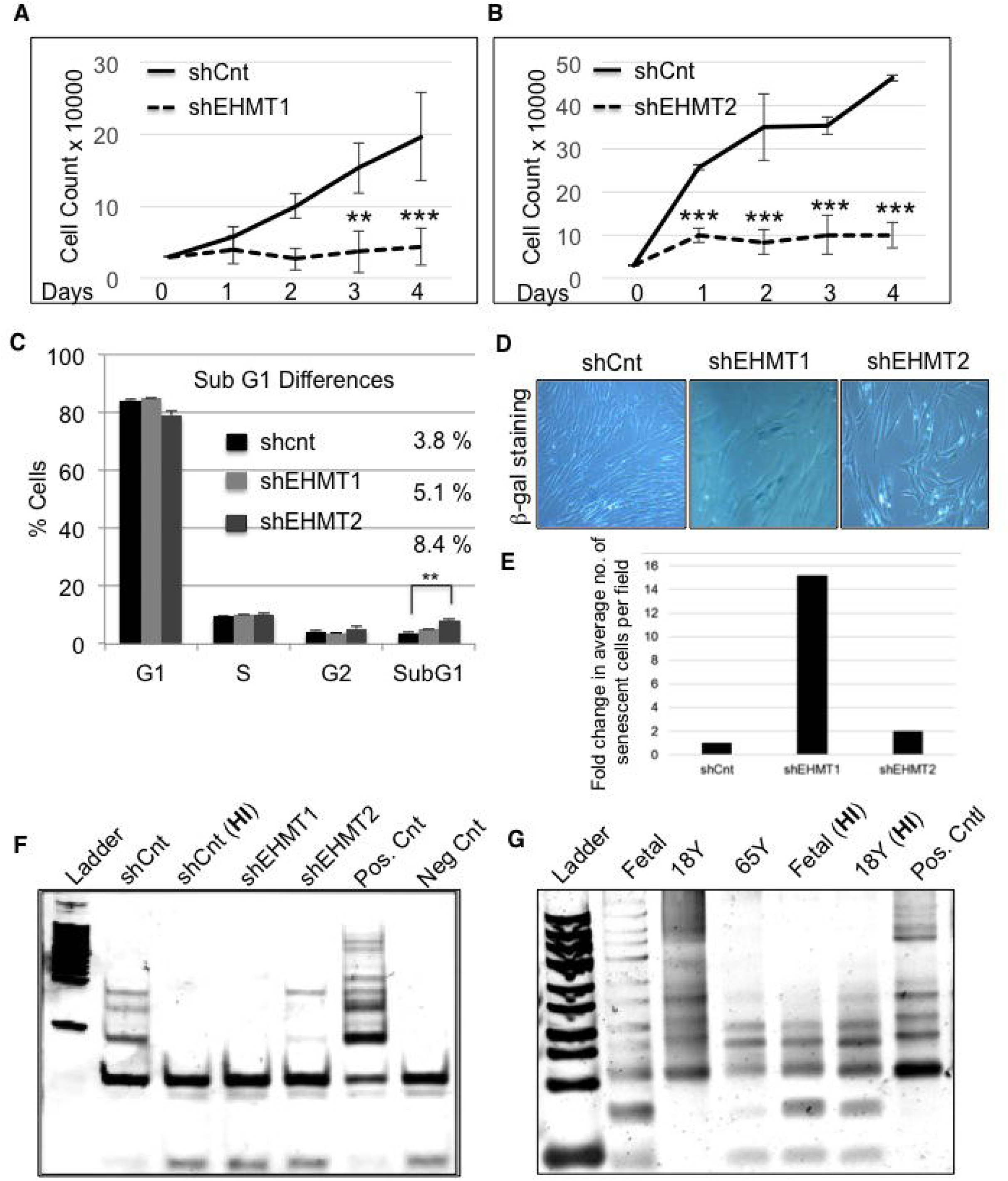
EHMT1 and EHMT2 knockdown leads to reduced cell proliferation and drives cells towards senescence. **A, B** Knockdown of EHMT1 and EHMT2 reduces cell proliferation. An equal number of HDFs were seeded and transduced with shCnt, shEHMT1 and shEHMT2 virus. Forty-eight hours post transduction, the number of cells in the culture was counted over a period of four days as indicated. (n=3), for shEHMT1, Day3: shCnt vs. shEHMT1 (**p=0.0015); Day4: shCnt vs. shEHMT1 (***p<0.0001); for shEHMT2, Day1: shCnt vs. shEHMT2 (***p=0.0002), Day2: shCnt vs. shEHMT2 (***p<0.0001), Day3: shCnt vs. shEHMT2 (***p<0.0001), Day4: shCnt vs. shEHMT2 (***p<0.0001). (Two-way ANOVA, post-hoc: Tukey’s multiple comparison test) **C** Quantification of cell cycle analysis. Increase in sub-G1 population an indication of cell apoptosis upon EHMT2 knockdown. For shCnt, (n=3), shEHMT1, (n=2) and shEHMT2, (n=2) SubG1: shCnt vs. shEHMT2 (**p=0.0011) (Two-way ANOVA, post-hoc: Tukey’s multiple comparison test). **D, E** Equal number of HDFs were seeded and transduced with shCnt, shEHMT1 and shEHMT2 virus. β-Galactosidase (senescence) assay was performed to monitor the cellular senescence in cultures. Fold change in an average number of senescent cells per field was quantitated. **F, G** TRAP assay to detect telomerase activity in indicated age groups as well as in shCnt, shEHMT1 and shEHMT2 transduced cells. Fetal, 18Y and shCnt fibroblast lysates were heat inactivated (HI) as a negative control for the assay. Telomerase activity was reduced upon knockdown of EHMT1 and EHMT2 as well as during physiological aging from fetal to 65Y old cells.

The telomerase enzyme prevents the replicative senescence in primary fibroblast. Examining the levels of telomerase activity in shEHMT and aged cells identified a correlation between the loss of EHMT proteins with reduced telomerase activity (Fig 7F and G). Overall our results demonstrated that the depletion of EHMT1 and EHMT2 was correlated with reduced proliferation.

## Discussion

The NL is a meshwork of lamins that constitutes the nucleoskeleton required for nuclear structure and function [8–10,31]. The NL undergoes extensive posttranslational modifications (PTMs) that are crucial for their localization to regulate a variety of biological processes [32]. While uncovering the mechanism by which EHMTs organize heterochromatin, we have identified lysine methylation as a novel PTM on the nuclear localization signal (NLS) of LMNB1 that is critical for its retention at the NP and maintaining NL stability. Interestingly, this NLS motif is conserved in LMNA/C and is required for lamin-chromatin interactions [33]. High resolution imaging of endogenous LMNA and LMNB1 demonstrated that individual homopolymers exist in close contact with each other [28]. Our results showing that aggregates of mutant-LMNB1 contain LMNA/C indicate the potential crosstalk between the two proteins via PTMs, thereby opening new avenues to explore the role of methylated LMNB1 towards the assembly of NL and its integrity. LMNA is extensively studied with several known binding partners and disease causing mutations [34]. However, patients with laminopathies rarely exhibit a defect in the NLS sequence likely due to its lethality. In this regard, our study offers a new perspective on the less studied LMNB1 in the context of normal physiology and perhaps in laminopathies/disease.

Methylation of lysine residues facilitates a variety of functions including protein stability [35–38]. Altogether our data demonstrated EHMT1 and EHMT2 as upstream regulators of LMNB1 that influences its protein levels via PTMs. While EHMT2 is known to methylate a variety of non-histone proteins [24], our study for the first time demonstrates the competency of EHMT1 to methylate non-histone proteins and utilizing it as a mechanism to attach heterochromatin to the NP during aging. Structurally similar EHMT1/2 proteins form a heteromeric complex in mammalian cells [23] and are known to fulfill both overlapping and unique physiological roles in developing and adult animals [39]. In the quest to understand the individual contributions of EHMT proteins in regulating peripheral heterochromatin, we identified that both EHMTs regulate LMNB1. However, unique molecular changes seen upon EHMT1 loss such as disruption of NE integrity coupled with the loss of architectural proteins like HP1 and HMG that influence heterochromatin organization requires further investigation. Nonetheless, our studies provide a broader role for EHMTs by which it impacts the spatial distribution of the genome.

There are a number of studies demonstrating redistribution or loss of chromatin modifiers and their implications in aging [40,41]. These studies mainly focused on the consequence of the global loss of chromatin structure but none addressed the mechanisms underlying the alteration of genome architecture. Our study not only demonstrates the correlation between the expressions of EHMTs with peripheral heterochromatin organization during aging but also provides a mechanism by which EHMT regulates higher-order chromatin structure via stabilization of the NL and architectural proteins. These results are supported by previous observations wherein defects in the sophisticated assembly of nuclear lamins along with architectural proteins results in disease or aging [41–45]. Reorganization of heterochromatin at the NP by restoration of EHMT1 or EHMT2 in aged cells further reinforces the fact that EHMT proteins are key determinants of higher-order chromatin organization.

Aging associated defects in chromatin organization exhibit a variety of functional consequences such as misregulation of gene expression via alteration of the epigenome, activation of repeat elements and susceptibility to DNA damage [46,47]. Additionally, loss of lamins leads to altered mechano-signaling [48]. Together, these processes make aged cells stressed and also influence the stress response contributing towards reduced proliferation and enhanced senescence. Our studies revealed a direct correlation between loss of peripheral heterochromatin and reduced proliferation in shEHMT and 65Y HDFs. Taken together, we conclude that the steady loss of EHMT proteins drives normal aging in fibroblasts. It remains a mystery as to how EHMT proteins are regulated. We propose that EHMT2 degradation leads to gradual destabilization of EHMT1 in response to aging via currently unknown mechanisms. It is important to note that these results are in cultured HDFs and yet to be established if this extends in the context of tissue aging.

## Materials and Methods

### Antibodies & Inhibitors

The following antibodies were used in the current study: EHMT1 (A301-642A, Bethyl Laboratories, Rabbit polyclonal), EHMT1 (NBP1-77400, Novus Biologicals, Rabbit polyclonal), EHMT2 (NBP2-13948, Novus Biologicals, Rabbit polyclonal), EHMT2 (07-551; Millipore, Rabbit polyclonal), H3K9me2 (ab1220, Abcam, Mouse monoclonal), LMNB1 (ab16048, Abcam, Rabbit polyclonal), LMNB2 (ab8983, Abcam, Mouse Monoclonal), LMNA/C (sc-20681, Santacruz, Rabbit polyclonal), HP1-β (ab101425, Abcam, Mouse monoclonal), H3K9me3 (ab8898, Abcam, Rabbit polyclonal), H3K27me3 (07-449, Millipore, Rabbit polyclonal), H3 (ab1791, Abcam, Rabbit polyclonal), GAPDH (G9545, Sigma, Rabbit polyclonal), Anti-Methyl lysine antibody (ICP0501, Immunechem, Rabbit polyclonal), Anti-6X His-tag antibody (ab9108, Abcam, Rabbit polyclonal), Anti-GST (ab9085, Abcam, Rabbit polyclonal), Anti-GFP (ab290, Abcam, Rabbit polyclonal), Ash2L (ab176334, Abcam, Rabbit monoclonal), p16 antiboy (ab54210, Abcam) Normal Rabbit IgG (12-370, Millipore), Normal Mouse IgG (12-371, Millipore). Secondary antibodies Alexa Fluor 488 Goat anti-mouse (A-11001), Alexa Fluor 488 Goat anti-rabbit (A-11008) and Alexa Fluor 568 Goat anti-rabbit (A-11011) were purchased from Thermo Fisher Scientific while secondary antibodies Goat anti-mouse HRP (172-1011) and Goat anti-rabbit HRP (170-8241) were from Bio-Rad. MG132 (C2211) and BIX 01294 (B9311) inhibitors were purchased from Sigma-Aldrich.

### Cell lines

Fetal (2300), Neonatal (2310) 18Y (2320) old HDFs were purchased from ScienCell, 38Y old HDFs (CC-2511) from Lonza while 19Y (C-013-5C), 31Y (C-013-5C), 37Y (CC-2511), 40Y (C-013-5C) and 65Y (A11634) old HDFs were purchased from Thermo Fisher Scientific. HDFs obtained from individuals of different age groups were controlled for gender (female) and ethnicity (Caucasian).

### shRNA constructs and Virus preparation

EHMT1 shRNA-V3LHS_36054, was purchased from Sigma. The vector was co-transfected with psPAX2, pMDG2 in 293-LX packaging cell line using lipofectamine LTX (15338500, Thermo Fisher Scientific). Viral supernatants were harvested 48 h post transfection and concentrated using Amicon Ultra-15 centrifugal filter units (UFC910024, Millipore). Retroviral pSMP-Ehmt1_4 (plasmid # 36338, Addgene), pSMP-EHMT2_4 (Plasmid # 36334, Addgene), pSMP-EHMT2_1 (Plasmid # 36395, Addgene) vectors were purchased from addgene. These vectors were transfected in AmphoPack-293 cell line (631505, Clontech). Viral supernatants were harvested 48 h post transfection. Viral supernatant was used to transduce HDFs. V5-EHMT1 and Flag-EHMT2 constructs were obtained from Dr. Marjorie Brand (OHRI, Ottawa, Canada).

### Transfection

HEK293 cells were maintained in Dulbecco’s Modified Eagle Medium (DMEM) (10566-016, Thermo Fisher Scientific) supplemented with 10% fetal bovine serum (FBS) (10082147, Thermo Fisher Scientific), 2mM L-glutamine (25030081, Thermo Fisher Scientific), and 1% nonessential amino acids (NEAA, 11140050, Thermo Fisher Scientific). At 70% confluency, HEK293 cells were transfected with pEGFPC1, pEGFP-ANK or pEGFP-SET plasmids using Lipofectamine 2000 (11668019, Thermo Fisher Scientific). HDFs were maintained in DMEM supplemented with 10% FBS, 2mM L-glutamine and 1%NEAA. For knockdown experiments, cells were transduced with viral particles containing shRNAs against EHMT1 or EHMT2 and incubated for 48 h. Post 48 h transduction, cells were washed with complete media. Transduced fibroblasts were further cultured for 48 h, harvested for protein and RNA extraction. For proteasome degradation pathway inhibition experiments, cells were treated with MG132 for 6 h at 10μM concentration. For BIX 01294 experiments, cells were treated at 1μM concentration for 48 h. Respective inhibitor treated cells were further processed for western blot, immunostaining &/or electron microscopy.

Transfections of Wt.LMNB1 and K417A-LMNB1 were carried out in fetal HDFs using Neon transfection method (MPK10096, Thermo Fisher Scientific). Briefly 5 × 10^5^ − 2 × 10^6^ cells were collected and the pellet was washed twice with PBS. Cell pellet was then resuspended in Buffer R along with 2-3 μg of respective plasmids. Cell suspension was electroporated using Neon pipette and immediately transferred to pre-warmed media. Transfected cells were seeded on coverslips and were processed for immunostaining or Electron microscopy 6 h post transfection. Images for Wt.LMNB1 and K417A-LMNB1 transfected cells were acquired at the same settings. Over expression of EHMT1 and EHMT2 was carried out by transfecting V5-EHMT1 and Flag-EHMT2 over expression constructs in old HDFs using Neon transfection method (MPK10096, Thermo Fisher Scientific) as described above. Transfected cells were seeded on coverslips and were processed for immunostaining or Electron microscopy 24 h post transfection.

### Cell growth curve

Fetal fibroblasts transduced with shCnt or shEHMT1 and shEHMT2, were independently seeded per well of a 6 well plate, with one well each for different time points. After each time point, cells were harvested and cell count was determined. Cell count was plotted against the time points to determine the growth curve.

### SA-β-Galactosidase Assay

SA-β-Galactosidase staining was performed using the Senescence Cells Histochemical Staining Kit (CS0030, Sigma). Cells were rinsed with PBS followed by fixing with 1X Fixation buffer provided with the kit for 8 min at RT. After rinsing thrice with PBS, 0.5ml of the staining mixture was added and incubated at 37°C without CO_2_ for 18 h. The percentage of β-gal positive cells were quantified from the images taken at 10 randomly selected microscopic fields.

### Cloning

The Ankyrin and SET domains of EHMT1 were amplified from cDNA prepared using the Superscript III cDNA synthesis kit (11752-050, Thermo Fisher Scientific) from HDFs with the help of Ankyrin (737-1004 AA) and SET (1013-1265 AA) domain primers (Appendix Table S1). The PCR amplified EHMT1-Ankyrin and SET products were then cloned into pEGFPC1 vector (6084-1, Clontech) to generate the plasmid constructs, pEGFP-ANK and pEGFP-SET. For cloning the C-Term of LMNB1, the cDNA from HDFs was PCR amplified using primers LMNB12F and LMNB1-1R (Appendix Table S1) and cloned into pET28a+ vector between BamHI and HindIII restriction digestion sites. The identity of all plasmids was confirmed by sequencing.

Site directed mutagenesis was carried out at 417^th^ lysine residue in LMNB1 plasmid construct mWasabi-LaminB1-10 (411^th^ position in mWasabi-LaminB1 plasmid and 417^th^ in Uniprot LMNB1 sequence). mWasabi-LaminB1-10 was a gift from Michael Davidson (Addgene plasmid # 56507). Briefly, 200 ng of the LMNB1 plasmid construct was subjected to a standard mutagenic PCR reaction with *Q5* High Fidelity DNA polymerase (M0491, New England Biolabs) and 25 ng of specific primers. The primers used.

~~~
EHMT1 ANK: F- CCGGAATTCTTC CAC CCA AAG CAG CTG TAC
           R- AAGGAAAAAAGCGGCCGCGCTGGGCCTGTCGGG
pEGFPC1-ANK Construct
EHMT1 SET: F- CCGGAATTCGGACATCGCTCGAGGCTACGAG
           R- AAGGAAAAAAGCGGCCGCGTGCCGGCACTTGGG
pEGFPC1-SET Construct
LMNB1-CT: F- TCGCGGATCC atgtatg aagaggagat taac
           R- GGGAAGCTT ttacataattgcacagcttcta
~~~

The mutagenic PCR reaction parameters were as follows: 98°C for 1 min, 18 cycles (98°C for 15 sec, 70°C for 15 sec, 72°C for 2 min) and 72°C for 5 min. The final reaction volume was 50 μL. The reaction product was digested with 10 U of methylation-sensitive enzyme *DpnI* at 37°C for 1h. (R0176, New England Biolabs). *E. coli* DH5-α competent cells were transformed with the amplified products. Finally, the plasmids were purified using the Qiagen plasmid DNA purification kit. Sequence for the K417A plasmid is provided in Supplementary Text 1.

~~~
Primers used for site directed mutagenesis
LMNB1-K417L: GAAAGCGGGCGAGGGTTGATGTG
LMNB1-K417L: AACCCTCGCCCGCTTTCCTCTAG
~~~

### qPCR and PCR

Total RNA was isolated using Trizol reagent (15596018, Thermo Fisher Scientific) as per manufacturer instructions. RNA was subjected to cDNA synthesis using Maxima First strand cDNA synthesis kit (K1641, Thermo Fisher Scientific). Semi-quantitative PCR reactions were performed using 2X PCR Master Mix (K0171, Thermo Fisher Scientific). Products were resolved on 1.2% agarose gels. Quantitative PCRs were performed using Maxima SYBR green qPCR master mix (2X) (K0251, Thermo Fisher Scientific). GAPDH was used as an internal control. Primer sequences used are

~~~
**TCF3** F- GTTTGAAACGGCGAGAAGAG R-CGGAGGCATACCTTTCACAT
**FGFR1:** F- GAACAGGCATGCAAGTGAGA R- GCTGTAGCCCTGAGGACAAG
**FOXM1:** F- TCCAGAGACTGCCAGAAGGT R-AAAAAGGACTCTGGCAAGCA
**CDK2B:** F-CGCCCACAACGACTTTATTT R-TTCGCTTCATGGTGAGTGTC
**LMNA:** F-AAGCAGCGTGAGTTTGAGAG R-TGTCCAGCTTGGCAGAATAAG
**LMNB1:** F- TTAGCCCTGGACATGGAAATC R- GATACTGTCACACGGGAAGAAG
**EZH2:** F- TCCTCCTGAATGTACCCCCA R- TGCACTTACGATGTAGGAAGC
**EHMT1:** F- GACATCAACATCCGAGACAA R- GAAAGAAAGAGGACGACACAG
**EHMT2:** F- GCCATGCCACAAAGTCATTC R- CTCAGTAGCCTCATAGCCAAAC
**GAPDH:** F-GAAATCCCATCACCAATCTTCCAGGR- GCAATTGAGCCCCAGCCTTCTC
~~~

### Preparation of whole cell extracts

HEK293 cells transfected with pEGFPC1, pEGFP-ANK and pEGFP-SET constructs were centrifuged and cell pellets were washed with ice-cold PBS and lysed in ice-cold RIPA buffer containing protease inhibitor cocktail (PIC) (11697498001, Sigma) per 10^6^ cells for 10 min on ice. The cell lysates were cleared by centrifugation and the supernatant was transferred to a fresh tube. Supernatant was used for IP experiments followed by western blotting.

### Protein Induction & purification from bacterial cells

The plasmids containing the human EHMT1-SET domain, EHMT1 (2IGQ) and EHMT2-SET domain, (Addgene plasmid # 25504 and Addgene plasmid # 25503 respectively) were expressed in *Escherichia coli C43 (DE3)* while LMNB1-CT was expressed in *Escherichia coli B834 (DE3)* cultured in LB medium with 50 μg/mL of kanamycin (Sigma). EHMT1 (2IGQ) (Addgene plasmid # 25504) and EHMT2 (Addgene plasmid # 25503) were a gift from Cheryl Arrowsmith. After induction, cells were harvested and resuspended in lysis buffer containing 25 mM Tris pH 8.0, 0.5 M NaCl, 2 mM L-Dithiothreitol (DTT, M109, Amresco), 5% glycerol, 0.05% Nonidet P-40 Substitute (M158, Amresco); 3 μg/ml of deoxyribonuclease I (DNase I, DN25, Sigma), 50 mM imidazole supplemented with Complete protease inhibitor cocktail and 0.2 mM phenyl methyl sulfonyl fluoride (PMSF, 93482, Sigma). Cells were lysed by sonication using Vibra Cell Sonicator (Sonics & Materials Inc.). The crude extract was cleared and the supernatant was incubated overnight with 2 ml Ni-NTA Agarose resin pre-washed with the lysis buffer described above. The resin was packed into Econo-Column (738-0014, Biorad). The column was washed and 6x His tagged protein was eluted from the resin in protein elution buffer supplemented with 250 mM and 500 mM imidazole. The purity of protein was assessed by SDS–PAGE. The purified protein was concentrated, buffer exchanged and protein dialysis was performed using Amicon Ultra-4 centrifugal concentrators (UFC801008, Millipore,) with a molecular weight cut off of 10 kDa and the final concentration was estimated using the Bradford protein assay (5000006, Bio-Rad). The protein was also subjected to mass spectrometry to assess its purity and molecular weight. In-gel digestion for mass-spectrometry analysis revealed a Mascot Score of 2354.46 for 6X His EHMT1-SET.

### Protein-protein interaction assays

For interaction assays, Ni-NTA beads pre-washed with IP100 buffer (25 mM Tris pH 7.5, 100 mM KCl, 5 mM MgCl_2_, 10% glycerol, 0.1% NP-40 and 200 μM PMSF) were incubated with 300 to 500 ng of 6X His-EHMT1-SET. The beads were washed twice with IP100 buffer followed by two washes with Flag buffer (20 mM HEPES, 150 mM KCl, 2 mM MgCl_2_, 0.2 mM EDTA pH 8.0, 15% glycerol, 0.1% NP-40 and 200 μM PMSF) both supplemented with 100 mM imidazole. The beads were then incubated with LMNB1-GST (H00004001-P01, Novus Biologicals) or GST alone (negative control) followed by washes with IP100 buffer and Flag buffer as mentioned above. The bead bound proteins were eluted and subjected to western blotting with anti-GST and anti-6x His antibodies.

### Methyltransferase assay

#### Detection by Western blot

LMNB1-CT and EHMT1-SET/ EHMT2-SET were incubated along with 50 μM SAM in methyltransferase assay buffer in a 50 μl reaction at RT for 1 h. 12 μl of 4x-SDS-PAGE loading dye was added to all the tubes to stop the reaction, samples were heated at 95°C for 8 min and resolved by 10% SDS-PAGE. The reaction was split into two, one was used for Western blot to probe with Anti-Methyl-K antibody and the other half was used to stain with Coomassie Blue stain (Coomassie Brilliant Blue R-250, Amresco).

#### Detection by fluorescence

The Methyltransferase assay was performed as per manufacturer’s instructions (ADI-907-032, Enzo Life Sciences). Sequences for LMNB1 peptides used for the assay in Fig 2E are LMNB1-RKR: SVRTTRGK**RKR**VDVEESEA; LMNB1-RAR: SVRTTRGK**RAR**VDVEESEA

### Immunoprecipitation (IP)

For IP experiments, cell or nuclear lysates (400 ug) prepared from HEK293 or HDFs to Dynabeads Protein A (10001D, Thermo Fisher Scientific) that were prebound with 2–3μg of indicated antibody and incubated overnight at 4°C. Beads were then washed and eluted in 2X loading dye. Eluted proteins were subjected to western blotting with indicated antibodies.

### Sequential IP

For sequential IP, nuclear extract was added to Dynabeads Protein A (10001D, Thermo Fisher Scientific) pre-bound with 3-4 μg of EHMT1, EHMT2, LMNB1 and Rabbit IgG antibodies overnight at 4°C. These beads were then washed with IP 100 and FLAG buffer. Primary IP elution was carried out with 0.1M glycine pH 2.5-3 for 10 min. The elution was stopped by Tris buffer pH 9.5. The elute from EHMT1, EHMT2, LMNB1 and IgG were collected and incubated overnight at 4°C with Dynabeads Protein A pre-bound with EHMT2, EHMT1, EHMT1 and IgG respectively. The beads were subsequently washed with IP100 and FLAG buffer. The elution for secondary IP was carried out in sample buffer at 95°C. Next, eluted products were probed with respective antibodies in western blot as mentioned in the Fig 1B.

### Chromatin Immunoprecipitation (ChIP)

ChIP was performed as described previously Brand *et al* 2008 [49] with some modifications. In Brief, fetal HDFs were cross-linked with 1% formaldehyde. Cells were lysed in buffer N containing DTT, PMSF and 0.3% NP40. After isolation of nuclei, chromatin fractionation was done using 0.4U of MNase (N5386, Sigma) at 37°C for 10 min. Reaction was stopped using MNase stop buffer without proteinase K. Simultaneously, antibodies against EHMT1, LMNB1 and Rabbit IgG were kept for binding with Dynabeads for 2 h at RT. After equilibration of beads, chromatin was added for pre-clearing. To antibody bound beads pre-cleared chromatin was added and kept for IP at 4^0^C overnight.

Next day, beads were washed and eluted at 65°C for 5 min. Eluted product was subjected to reverse cross-linking along with input samples, first with RNAse A at 65°C overnight and then with proteinase K at 42°C for 2 h. After reverse cross-linking, DNA purification was performed using phenol-chloroform-isoamyl alcohol extraction method. The amount of DNA was quantitated using Qubit fluorometer (Thermo Fisher Scientific).

### ChIP-seq library preparation

ChIP DNA was subjected to library preparation using TruSeq ChIP sample preparation kit from Illumina (IP-202-1012). Briefly, ChIP samples were processed for end repair to generate blunt ends using end repair mix. A single ‘A’ nucleotide was added to the 3’ ends of the blunt fragments to prevent them from ligating to one another during the adapter ligation reaction. In the next step, indexing adapters were ligated to the ends of the DNA fragments. The ligated products were purified on a 2% agarose gel and narrow 250-300bp size range of DNA fragments were selected for ChIP library construction appropriate for cluster generation. In the last step, DNA fragments with adapter molecules on both ends were enriched using PCR. To verify the size and quality of library, QC was done on high sensitivity bioanalyzer chips from Agilent and the concentration was measured using Qubit dsDNA HS assay kit (Q32851, Thermo Fisher Scientific). After passing QC, samples were sequenced 75 paired end (PE) on NextSeq Illumina platform. Genotypic Technology Pvt. LTD. Bengaluru, India, performed the sequencing.

### ChIP-seq Analyses

Alignment of ChIP-seq derived short reads to the human reference genome (UCSC hg19) was done using Bowtie2 short read aligner [50] with default parameters. Subsequently, aligned ChIP-seq reads from two replicates were merged. Peak calling was done for each sample with their respective control using MACS 1.4 algorithm [51]. The following parameters deviated from their default value: - effective genome size = 2.70e+09, bandwidth = 300, model fold = 10, 30, p-value cutoff = 1.00e-03.

In order to identify regions enriched for EHMT1 and LMNB1, we employed a two-step approach, first total peak counts for EHMT1 and LMNB1 was calculated in 1MB window for all the chromosomes. Next, ratio of total peak count over expected peak count (total peaks from a chromosome divided by total 1 MB window for the same chromosome) was calculated for each 1MB window. Raw data has been deposited in NCBI {SRP110335 (PRJNA391761)}.

### RNA-seq and data analysis

Fetal HDFs transduced with shEHMT1, shEHMT2 or shCnt were harvested and RNA was extracted by Trizol method. RNA concentrations were estimated using Qubit fluorometer and quality was assessed using Bioanalyzer. After passing the QC, samples were subjected for library preparation and QC for the same. Samples were sequenced at Genotypic Technology Pvt. LTD. Bengaluru, India.

We obtained ~30-45 million reads from EHMT1 and EHMT2 knockdown samples. From the sequencing reads, adapters were trimmed using Trimmomatic program [52]. The reference based transcriptome assembly algorithms TopHat v2.1.0 [53], Cufflinks v2.2.1 [54] pipeline were used to assemble the transcripts with hg19 genome/transcriptome (http://genome.ucsc.edu/) [55] as reference annotation to guide RABT assembly. Cuffdiff v2.2.1 [56] was used to identify differentially expressed genes. The transcripts with adjusted *p* value < 0.05 & fold change > 1.5 were considered to be significantly expressed. We used GSEA [57] to identify top 100 pathways (FDR q value < 0.05).

To shortlist genes involved in epigenetic modifications, Epigenetic Modifiers (http://crdd.osdd.net/raghava/dbem/) [58] & Epifactors (http://epifactors.autosome.ru/description) [59] database were used. The aging related genes were shortlisted from GenAge (http://genomics.senescence.info/genes/allgenes.php) database [60]. Significantly expressed genes from both EHMT1 & EHMT2 knockdown datasets were overlapped with above mentioned databases. We used customized Perl scripts for all the analysis done in this study. All the plots and statistical analysis were done using R studio (R Development Core Team, 2011) [61]. Raw data has been deposited in NCBI {SRP110335 (PRJNA391761)}.

### Immunostaining

Briefly, cells were fixed with 4% Paraformaldehyde (PFA, P6148, Sigma) for 10 min at room temperature (RT) and permeabilized with 0.5% triton X-100. The blocking was done with 5% BSA for 1 h. Antibodies mentioned in the antibodies and inhibitors section were used at desired dilution and imaging was carried out on FV3000 Confocal Microscope (Olympus). Image analysis and extraction of raw files was done with the Fiji software [62].

### Transmission Electron Microscopy (TEM)

For TEM sample preparation cells of different age groups or treated with different conditions as mentioned in results sections were trypsinized and the pellet was fixed with 2.5% glutaraldehyde and 2% sucrose at RT for 1 h. Next, fixative was removed and pellet was washed with 0.1M phosphate buffer pH7.4. The buffer was replaced with 1% osmium tetraoxide and 1.5% potassium ferrocyanide and kept at 4°C for 75 min. Cell pellet was washed with 0.1M phosphate buffer as well as distilled water, subsequently dehydrated with gradient of ethanol followed by two changes of propylene oxide. Cell pellet was embedded gradually in Epoxy812 resin mixture (13940, EMS). Resin embedded pellets were allowed to polymerize at 60°C for 72 h. The blocks were trimmed; sections of 60nm size were collected and imaged with the Tecnai G2 Spirit Bio-TWIN Transmission Electron Microscope at National Centre for Biological Sciences (NCBS) EM facility.

### Flow cytometry

Cells were harvested and fixed in ice cold 70% ethanol for 30 min at 4°C. Post fixation, cells were pelleted, washed twice in PBS and resuspended in 0.5ml PBS containing 0.2mg/ml RNase A (EN0531, Thermo Fisher scientific) and incubated for 30 min at 37°C. To this Propidium Iodide (P3566, Thermo Fisher Scientific) at a concentration of 50 μg/ml was added and incubated for 10 min at 37°C. The percentage of cells in various phases of cell-cycle was assessed by flow cytometry (BD FACSVerse) and analyzed using the FlowJo software.

### Telomerase assay

Telomerase activity was detected using the PCR-based Telomeric Repeat Amplification Protocol (TRAP) assay kit (Millipore, S7700). Briefly, different age groups HDFs or cells transduced with shEHMT1 and shEHMT2 were harvested and resuspended in 1X CHAP Buffer provided with the kit. The protocol was followed as per the manufacturer’s instructions.

### Quantitation for mean fluorescence intensity

For quantitation of fluorescence intensity in Fig 4A, Supplementary Figure 2E, 4E, 4M and Supplementary Fig 7A-D, MATLAB programming was used. Briefly, centroid of nuclei was determined and 200-line scans from center to periphery of the cells were drawn. Each line was further divided into 200 points. Average intensity distribution was calculated for each nucleus by calculating mean of 200-line scans from center to periphery. Mean fluorescence intensity (MFI) was plotted for center vs. periphery of nuclei. Mean intensity profile with standard deviation for all the measured nuclei. The Script for MATLAB program is provided in Computer Code1.

### Statistics

The detailed statistical analysis and methods have been described in the figure legends along with the p-values for respective data sets. The data is represented as mean ± SD. For statistical analysis, GraphPad Prism version 7 software was used.

## Supporting information

Differential expression of shEHMT1

Differential expression of shEHMT2 genes

GSEA upregulated pathways for EHMT1

Chromatin modifiers for shEHMT1 and shEHMT2_Supplementary Table 4.xls

Aging related genes for shEHMT1 and shEHMT2

Rao and Ketkar et al LMNB1 mutant sequence-Supplementary Text 1.pdf

Rao and Ketkar et al_script_Computer Code 1.pdf

## Acknowledgments

We thank Drs. Colin Jamora, Arjun Guha and Pavan Kumar P for the helpful suggestions and critical reading of the manuscript. We thank members of Jamora lab and Sara Ripamonti for experimental help. We thank Ashish Dhayani for help with TEM imaging. We thank Febina Ravindran for discussions and insights. The authors acknowledge the facilities, and the scientific and technical assistance of the electron microscopy facility at National Centre for Biological Sciences (NCBS) -TIFR Bangalore. RNA sequencing was performed at the Genotypic Technology Private Limited, Bangalore, India. Confocal microscopy and Flow cytometry were performed at the Central Imaging and Flow Facility (NCBS/inStem). The authors acknowledge the facilities, and the scientific and technical assistance of the Electron Microscopy Facility at NCBS. This work was supported by grant from the Wellcome Trust/DBT India Alliance (to S.R.) and inStem core funding. A.K. is Career Development Fellow (CDF) supported by funds from NCBS. R.A.R is supported by Indian Council of Medical Research (ICMR)-Senior Research Fellowship (SRF). S.R. C.P and D.P. are Wellcome Trust/ DBT India Alliance Intermediate Fellows.

## Author contributions

R.R. performed experiments on lentivirus, retrovirus production, shRNA transduction, overexpression studies, PCRs, immunostaining, TRAP assays, IPs, cell growth and cell-cycle studies. A.K performed experiments on immunostaining, western blotting, electron microscopy, IPs, ChIP, PCRs, MG132 and BIX 01294 studies and statistical analysis. N.K. is responsible for cloning and purification of EHMT1 and LMNB1 truncation proteins and LMNB1 methylation studies. P.K. is responsible for analysis of ChIP Seq data. A.M. cloned mammalian EHMT1 truncations; performed IP and western blot analysis. V.L is responsible for bioinformatics analysis of RNA-Seq data sets. S.D and S.R. wrote program-using MATLAB for the analysis of immunostaining experiments. VKK performed western blotting, statistical analysis and methyltransferase assays. A.G provided intellectual inputs. D.P. supervised bioinformatics analysis of RNA and ChIP sequencing and provided intellectual inputs on the manuscript. M.B and C.P provided EHMT1 and EHMT2 overexpression constructs. S.R. conceptualized the project, designed the experiments and wrote the manuscript.

## Supplementary Figure Legends

**Supplementary Figure 1.**
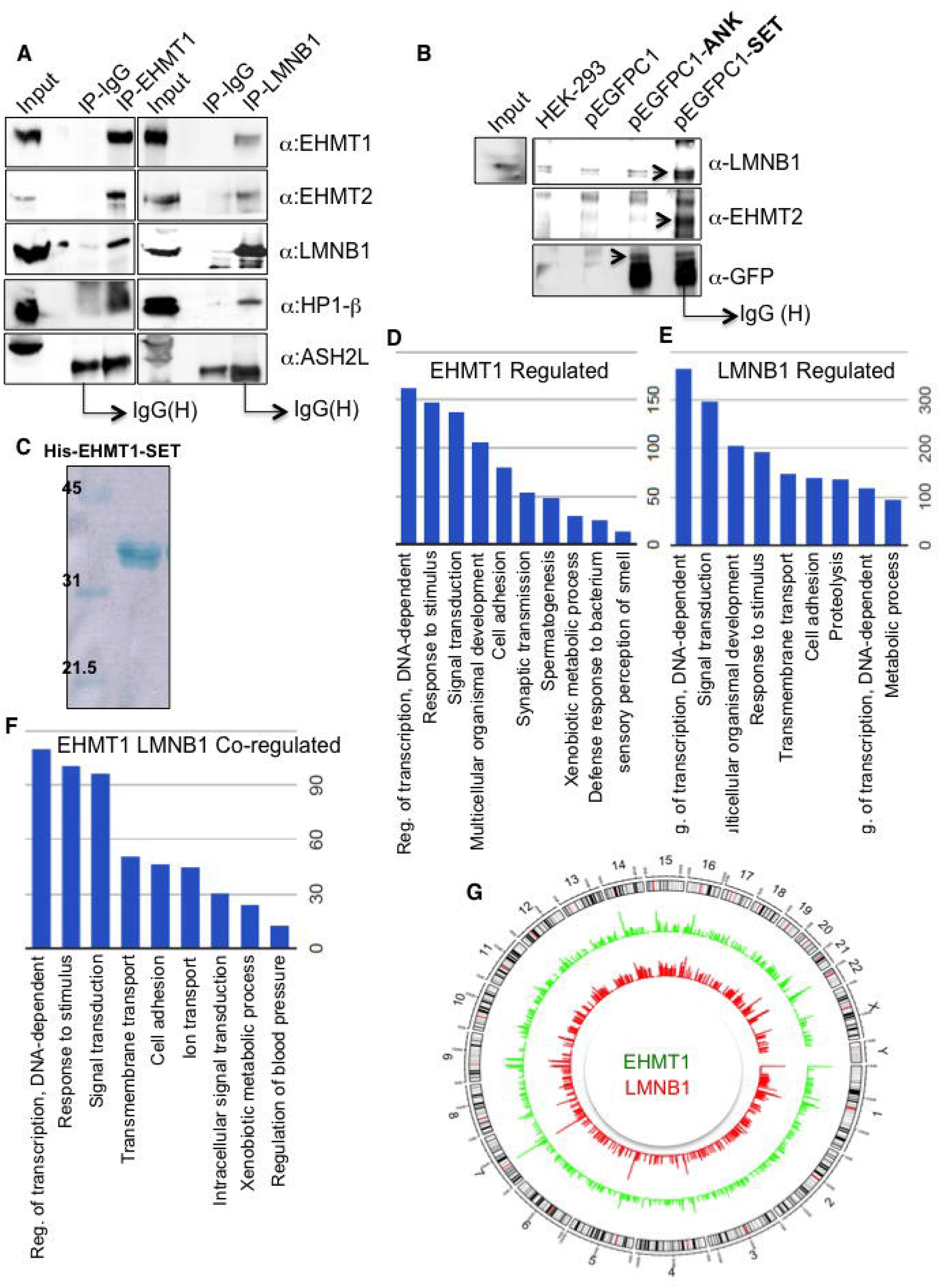
EHMT1 interacts with LMNB1 domain and co-regulate a subset of genes. **A** EHMT1 interacts with EHMT2, LMNB1, and HP1. Whole cell lysates from fetal HDFs were subjected to IP reaction using EHMT1 or LMNB1 antibody. Subsequently, western blot was performed using the IPed material to detect EHMT1, EHMT2, LMNB1, and HP1. ASH2L did not show any interaction with EHMT1 and LMNB1 thus acted as a negative control. **B** EHMT1 interacts with LMNB1 and EHMT2 via SET domain. HEK293 cells were transfected with pEGFP-Ankyrin (pEGFPC1-ANK) or pEGFPC1-SET domains of EHMT1 to determine domain specific association with LMNB1. Cell extracts were subjected to IP using GFP-antibody and the bound complexes were then analyzed by immunoblotting using LMNB1, EHMT2, and GFP antibodies. HEK-293 whole cell extract represents 1% input. pEGFPC1 empty vector transfected HEK293 or untransfected HEK293 cells were used as control reactions. Arrows indicate specific band. **C** Coomassie stained SDS-PAGE gel showing 6X His EHMT1-SET purified protein used for methyltransferase assays. **D, E, F** Gene Ontology (GO) analysis of EHMT1 and LMNB1 bound genes. Representative figure showing enriched GO terms for EHMT1 (**D**), LMNB1 (**E**) and EHMT1 and LMNB1 (**F**) co-bound genes. The length of the bar (y-axis) denotes total genes falling within GO term. **G** Circos plot showing genome wide peak density of EHMT1 (green) and LMNB1 (red).

**Supplementary Figure 2.**
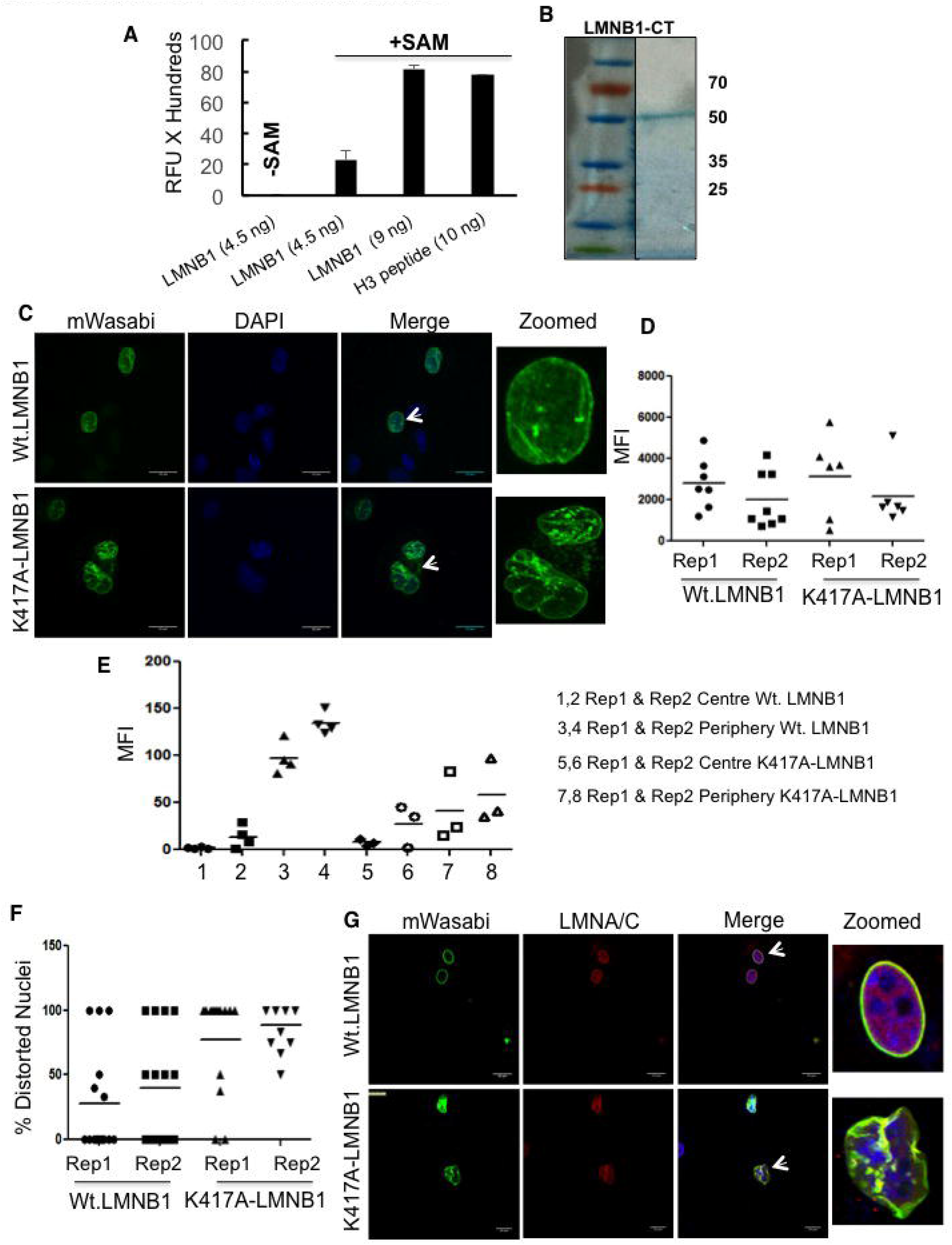
Mutation in LMNB1 causes distortion of the nuclear architecture. **A** Increasing concentrations of LMNB1-GST showed a greater degree of methylation by EHMT1-SET. Methyltransferase assay was performed using a fixed concentration of recombinant 6X His EHMT1-SET as an enzyme source, and SAM as a methyl group donor. Recombinant GST-LMNB1 (4.5 ng and 9 ng) and Histone H3 peptide (10 ng) were used as substrates in the assay. Histone H3 peptide was used as a positive control. The mean relative fluorescence unit (RFU) values represented in the graph were obtained after subtracting the values for controls EHMT1-SET only, SAM only, LMNB1-GST (4.5 ng) with those of the LMNB1-GST (4.5 ng and 9 ng) and Histone H3 (10 ng). **B** Coomassie stained SDS-PAGE gel showing 6X His LMNB1-CT purified protein used for methyltransferase assays. **C** mWasabi expression in fetal HDFs transduced with Wt.LMNB1 and K417A-LMNB1 mutant construct. (Scale bar: 20μm). Arrows indicate the cells zoomed in the far-right image presented. **D** MFI for mWasabi expression in cells transfected with Wt.LMNB1 and K417A-LMNB1 mutant constructs. Each biological replicate is represented as spread of technical replicates around the mean value. **E** MFI for mWasabi expression in cells transfected with Wt.LMNB1 and K417A-LMNB1 mutant constructs. MFI has been represented as center vs periphery of the nuclei. Individual biological replicates are plotted with the mean values. **F** Quantitation for percentage distorted nuclei in cells transfected with K417A-LMNB1 mutant construct compared to Wt.LMNB1 construct. Individual biological replicates are plotted with the median. **G** Immunostaining for LMNA/C in fetal HDFs transduced with Wt.LMNB1 and K417A-LMNB1 mutant construct. (Scale bar: 20μm). Arrows indicate the cells zoomed in the far-right image presented.

**Supplementary Figure 3.**
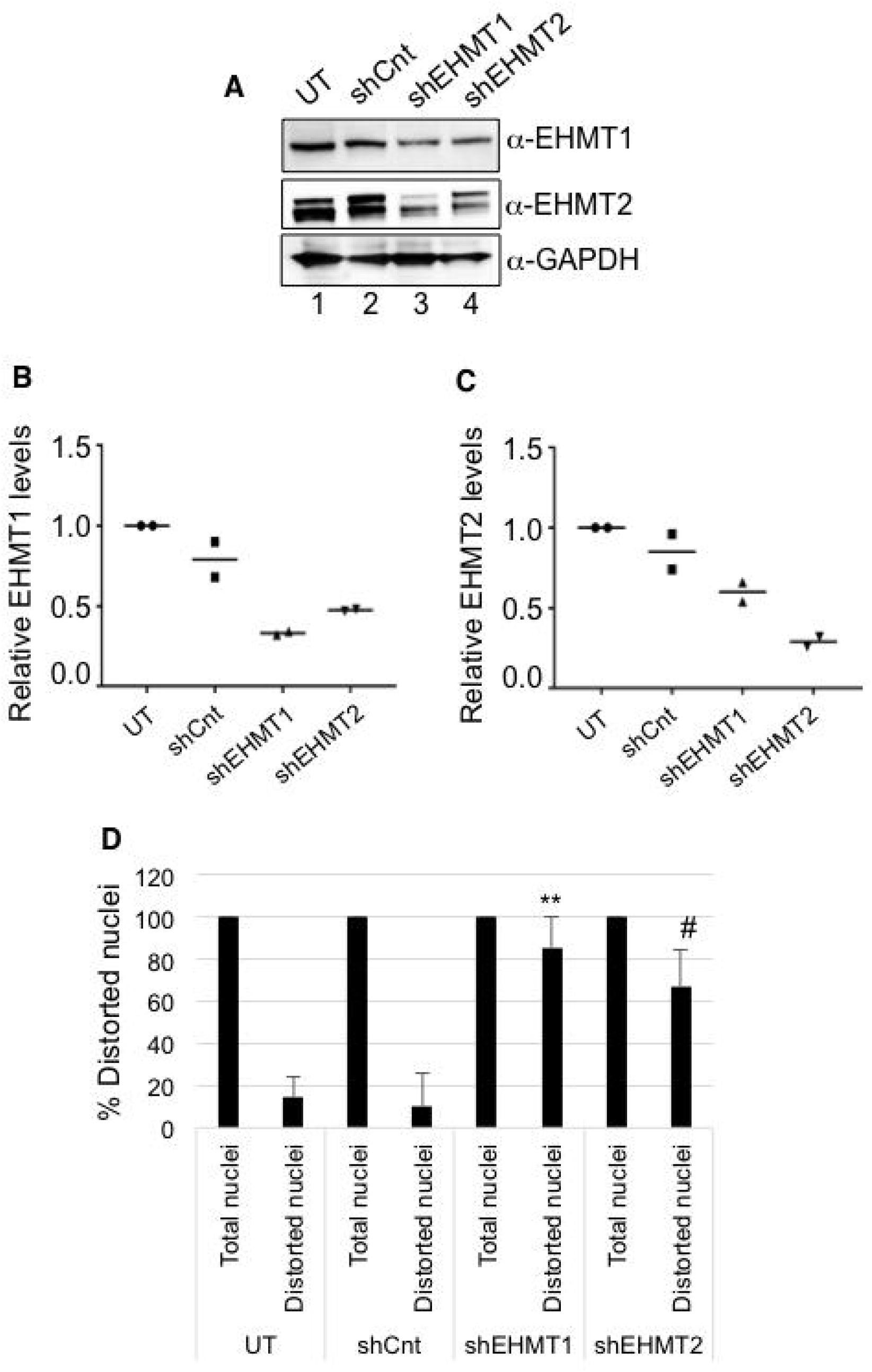
EHMT1 and EHMT2 knock-down causes nuclear distortion. **A** Western blot analysis for EHMT1 and EHMT2 in fetal HDFs upon knock down of EHMT1 and EHMT2. GAPDH was used as a loading control. **B, C** Relative protein levels of EHMT1 **(B)** and EHMT2 **(C)** upon knockdown. Each biological replicate is represented as spread of technical replicates around the mean value. **D** Percentage of distorted nuclei increases in human fibroblasts transduced with shEHMT1 and shEHMT2 virus compared to UT or shCnt. (n=3), For UT, (n=43); shCnt, (n=23); shEHMT1, (n=18); shEHMT2, (n=20) nuclei counted. shCnt vs shEHMT1, **p=0.0048; shCnt vs shEHMT2, #p=0.0153. (Kruskal-Wallis test, post-hoc test: Dunn’s multiple comparison test)

**Supplementary Figure 4.**
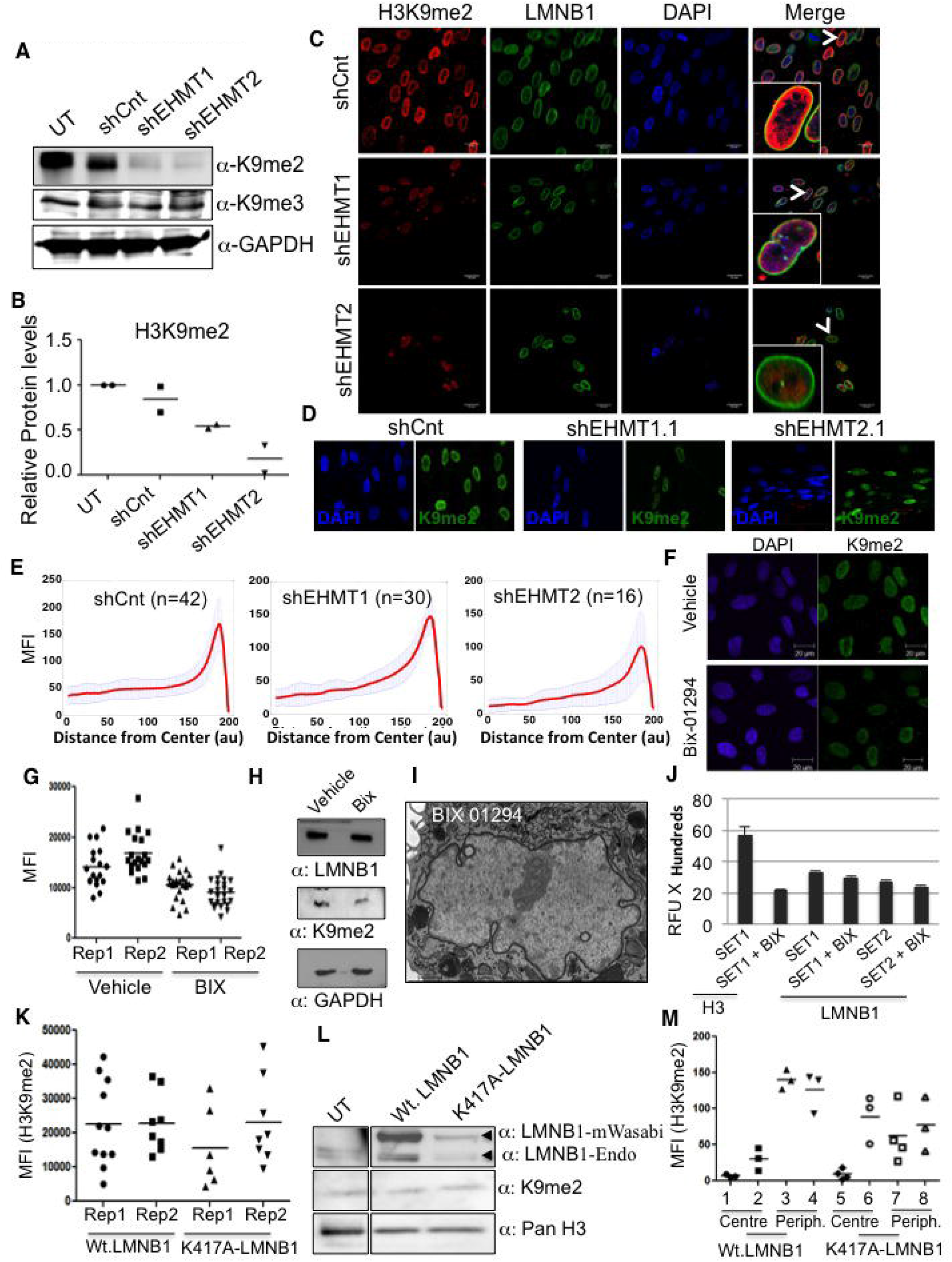
Knock-down of EHMT1 and EHMT2 reduces H3K9me2 marks. **A** Western blot analysis for H3K9me2 and H3K9me3 in UT, shCnt, shEHMT1 and shEHMT2 transduced HDFs. GAPDH was used as loading control. **B** Quantification of H3K9me2 upon EHMT1 and EHMT2 knockdown. Each biological replicate is represented as spread of technical replicates around the mean value. **C** Immunostaining for H3K9me2 co-stained for nuclear lamina using LMNB1 antibody in fetal HDFs transduced with shCnt, shEHMT1 and shEHMT2 virus. (Scale bar: 20μm). Insets are zoomed images of cells in the merge. **D** H3K9me2 immunostaining in fetal HDFs transduced with shEHMT1.1 (lentivirus) and shEHMT2.1 (retrovirus). (Scale bar: 20μm). **E** MFI profile (Centre to periphery) with standard deviation for all the measured nuclei of shCnt, shEHMT1 and shEHMT2. **F** Immunostaining for H3K9me2 in fetal HDFs treated with or without BIX 01294 (1μM) for 48 h. (Scale bar: 20μm). **G** MFI plot for H3K9me2 staining in fetal HDFs treated with or without BIX 01294 (1μM) for 48 h. Each biological replicate is represented as spread of technical replicates around the mean value. **H** Western blot for LMNB1 and H3K9me2 on cells treated with BIX 01294 or vehicle control. BIX 01294 treatment resulted in a reduction of H3K9me2 levels without altering LMNB1 expression. **I** *In vitro* methylation assay was performed using recombinant LMNB1-CT and EHMT1-SET or EHMT2-SET domain in the presence or absence of BIX 01294. H3 peptide was used as a positive control. **J** TEM image for fetal HDFs treated with BIX 01294. BIX 01294 treatment did not affect the peripheral heterochromatin distribution. (Scale bar: 1μm) **K** MFI for H3K9me2 staining in cells transfected with Wt.LMNB1 and K417A-LMNB1 constructs. Each biological replicate is represented as spread of technical replicates around the mean value. **L** Western blot showing expression of endogenous LMNB1 (top panel lower band) and Wasabi tagged LMNB1 (top panel upper band). The lysates from cells transfected with Wt.LMNB1 and K417A-LMNBI were probed with the LMNB1 antibody. The lysate was also probed with H3K9me2 antibody and H3 (loading control). **M** MFI for H3K9me2 staining in cells transfected with Wt.LMNB1 and K417A-LMNB1 constructs. MFI has been represented as center vs periphery of the nuclei. Each biological replicate is represented as spread of technical replicates around the mean value.

**Supplementary Figure 5.**
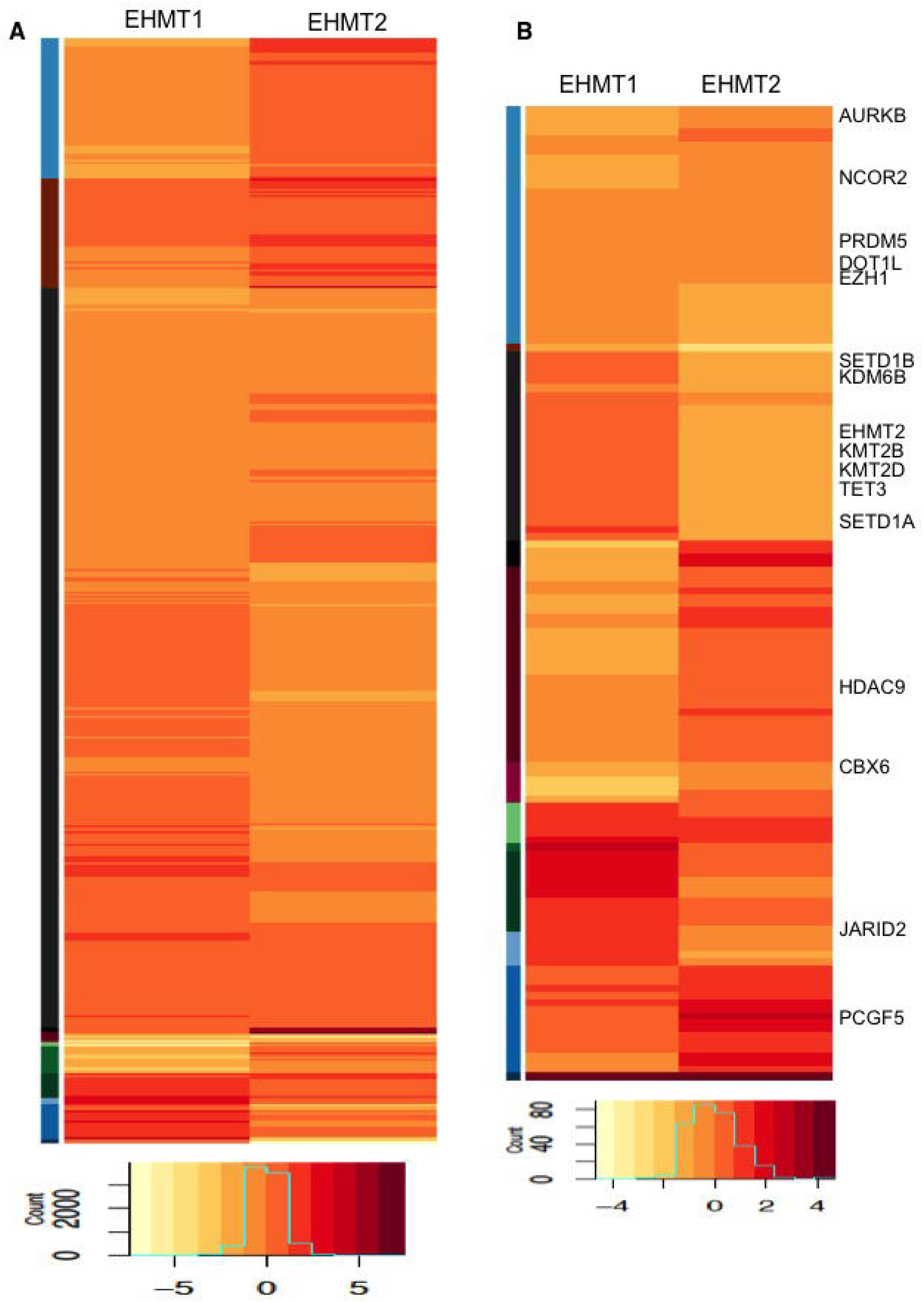
Transcriptome analysis of EHMT depleted cells. **A, B** Heat map for differential expression of **(A)** all genes and **(B)** chromatin modifiers obtained from RNA-seq analysis of shEHMT1 or shEHMT2 compared to shCnt transduced HDFs. Representative genes that were altered similarly or distinctly in EHMT1 vs EHMT2 are indicated from few clusters.

**Supplementary Figure 6.**
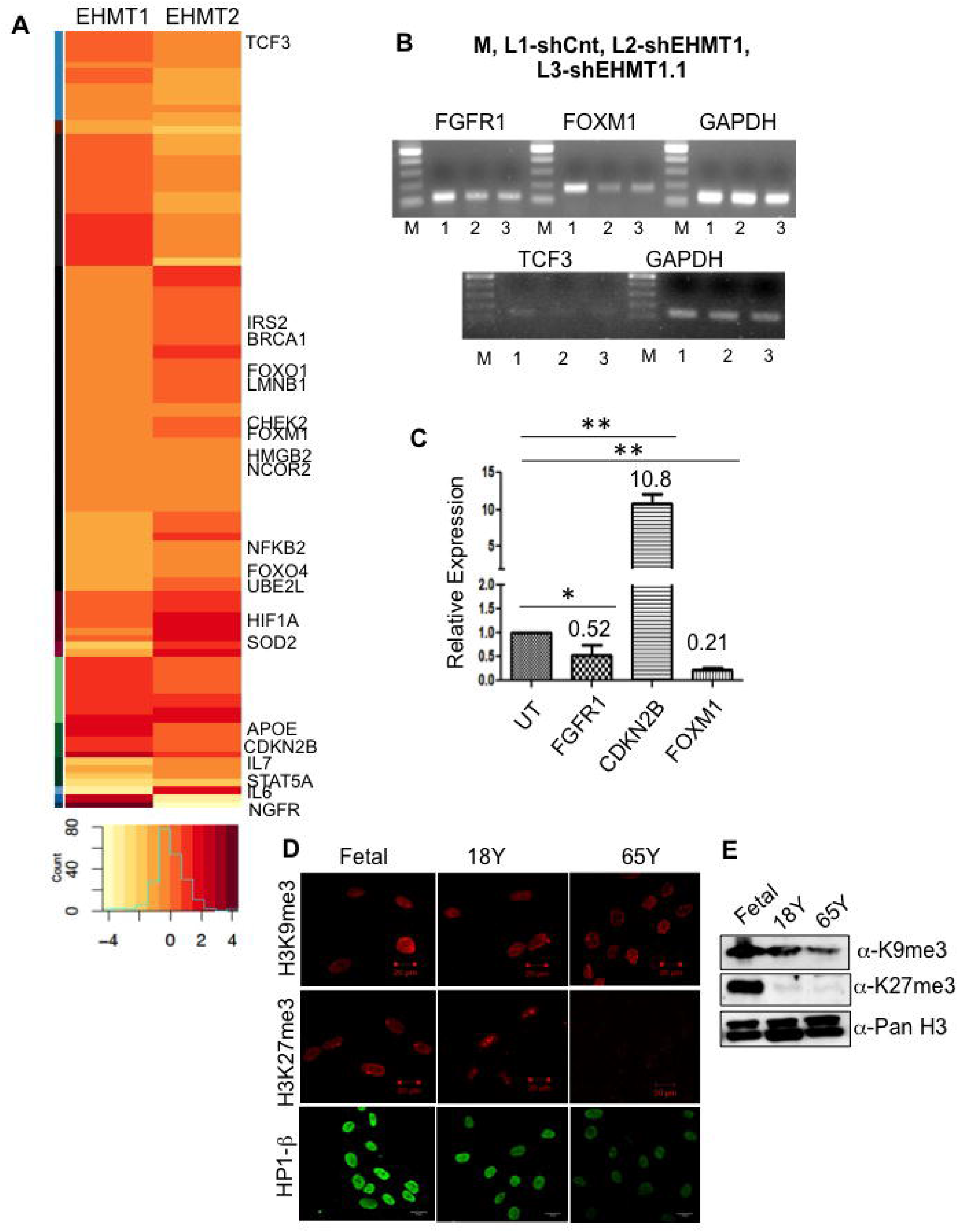
Depletion of EHMTs alters the expression of longevity-associated genes. **A** Heat map demonstrating differential expression of age related genes in EHMT1 and EHMT2 depleted fibroblasts. Representative genes that were altered similarly or distinctly in EHMT1 vs EHMT2 are indicated from few clusters. **B.** Validation for differential expression of candidate genes obtained from RNA-Seq analysis by semi-quantitative PCR. **C.** Validation for differential expression of candidate genes obtained from RNA-Seq analysis by qRT-PCR (n=3), UT vs FGFR1, p=0.05; UT vs CDKN2B, p= 0.0058; UT vs FOXM1, p=0.0018 One sample t-test, two tailed. **D.** Immunostaining for H3K9me3, H3K27me3 and HP1 in fetal, 18Y old and 65Y old HDFs followed by confocal imaging. (Scale bar: 20μm) **E.** Western blot analysis for H3K9me3 and H3K27me3 in fetal, 18Y and 65Y old HDFs.

**Supplementary Figure 7.**
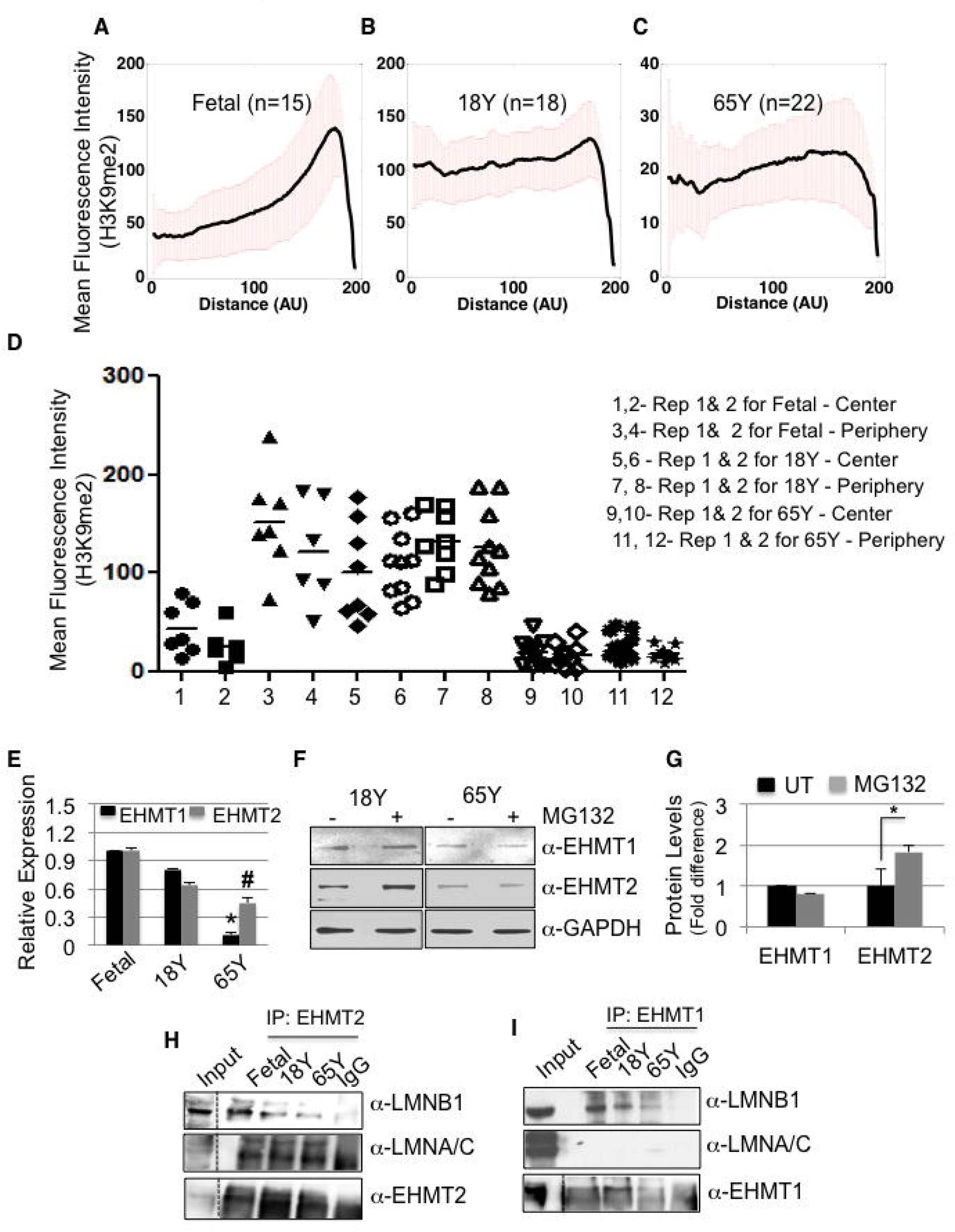
Interaction of EHMTs with LMNB1 during aging. **A-C.** MFI (Centre to periphery) with standard deviation for H3K9me2 staining in all the measured nuclei of fetal, 18Y and 65Y old HDFs. **D.** MFI for H3K9me2 staining in fetal, 18Y and 65Y old HDFs. MFI has been represented as center vs periphery. Individual biological replicates are plotted with mean values. **E** Relative expression of EHMT1 and EHMT2 at the mRNA level in fetal, 18Y and 65Y old HDFs. (n=3), For EHMT1: Fetal vs 65Y, *p=0.0219; For EHMT2: Fetal vs 65Y, #p=0.0219. (Kruskal-Wallis test, post-hoc test: Dunn’s multiple comparison test). **F** Western blot for EHMT1 and EHMT2 in 18Y and 65Y old HDFs treated with or without proteasomal degradation inhibitor MG132 (10μM for 6 h). **G** Quantification of EHMT1 and EHMT2 protein levels in 18Y old HDFs treated with or without MG132 treatment. (n=3), for 18Y EHMT1: UT vs MG132, p=0.5989, ns; For 18Y EHMT2: UT vs. MG132 (*p=0.0154). (One sample t-test, two-tailed). **H-I.** Cell lysates prepared from human fibroblasts of indicated age groups were subjected for IP using EHMT1/EHMT2 antibody. IPed material was analyzed by immunoblotting using LMNB1 and LMNA/C antibodies. 30 μg of fetal cell lysate was used as input control. Dotted lines indicate that different exposures were used for Input and IP of the same western blot.

**Supplementary Figure 8.**
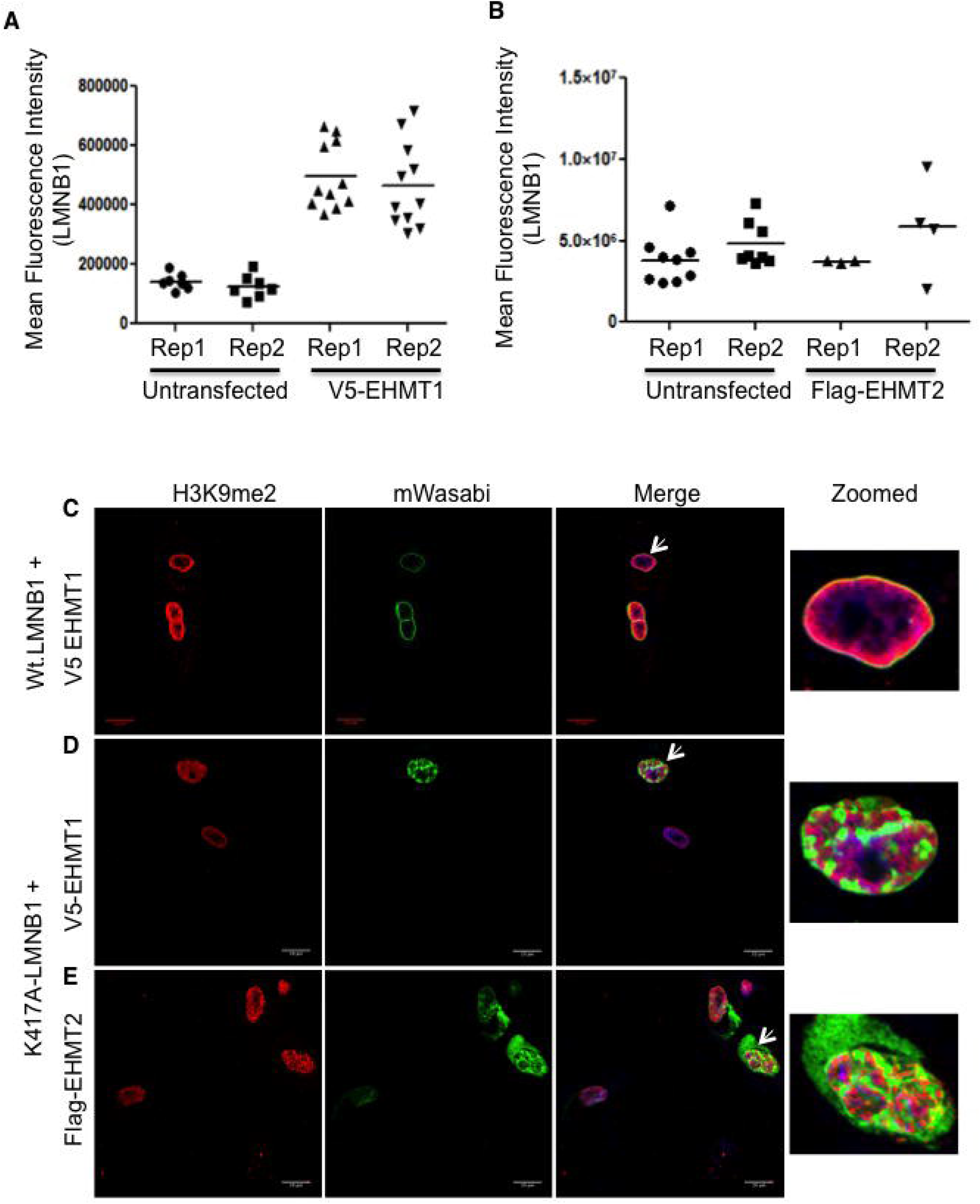
Co-expression of mutant LMNB1 with EHMTs aggregates H3K9me2 in the nucleoplasm. **A-B.** Quantitation of LMNB1 expression in aged HDFs upon overexpression of EHMT1 and EHMT2. Overexpression of EHMT1 significantly increased the LMNB1 expression while EHMT2 transfected cells did not show any change compared to untransfected cells. Biological replicates of V5-EHMT1 and Flag-EHMT2 expressing cells were plotted with the mean values. **C, D,E.** Old HDFs expressing Wt. LMNB1 + V5-EHMT1, K417A-LMNB1 + V5-EHMT1 and K417A-LMNB1 + Flag-EHMT2 were stained with H3K9me2 antibody. Mutation at lysine 417 position of LMNB1 affects the overall distribution H3K9me2 and morphology (Scale bar: 20μm). Arrows indicate the cells zoomed in the far right image presented.

**Supplementary Figure 9.**
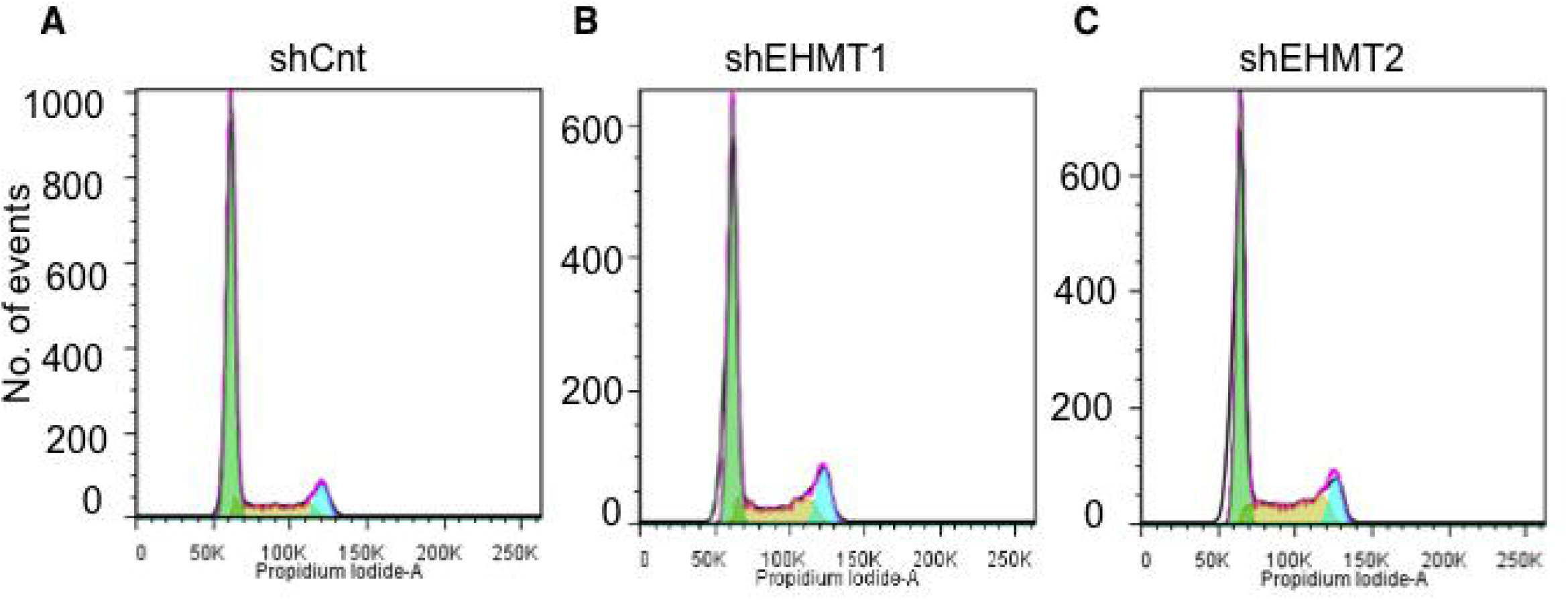
Physiological aging results in cell cycle arrest. **A-C** Cell cycle analysis for shCnt, shEHMT1 and shEHMT2 transduced fetal HDFs.

