## Supplementary material for "KMT1/Suv39 methyltransferase family regulates peripheral heterochromatin tethering via histone and non-histone protein methylations": Rao and Ketkar et al LMNB1 mutant sequence-Supplementary Text 1.pdf

### Wt. LMNB1Sequence

ATGGCGACTGCGACCCCCGTGCCGCCGCGGATGGGCAGCCGCGCTGGCGGCCCCACCACGCCGCTGAGC
C
CCACGCGCCTGTCGCGGCTCCAGGAGAAGGAGGAGCTGCGCGAGCTCAATGACCGGCTGGCGGTGTACA
T
CGACAAGGTGCGCAGCCTGGAGACGGAGAACAGCGCGCTGCAGCTGCAGGTGACGGAGCGCGAGGAGGT
G
CGCGGCCGTGAGCTCACCGGCCTCAAGGCGCTCTACGAGACCGAGCTGGCCGACGCGCGACGCGCGCTC
G
ACGACACGGCCCGCGAGCGCGCCAAGCTGCAGATCGAGCTGGGCAAGTGCAAGGCGGAACACGACCAGC
T
GCTCCTCAACTATGCTAAGAAGGAATCTGATCTTAATGGCGCCCAGATCAAGCTTCGAGAATATGAAGC
A
GCACTGAATTCGAAAGATGCAGCTCTTGCTACTGCACTTGGTGACAAAAAAAGTTTAGAGGGAGATTTG
G
AGGATCTGAAGGATCAGATTGCCCAGTTGGAAGCCTCCTTAGCTGCAGCCAAAAAACAGTTAGCAGATG
A
AACTTTACTTAAAGTAGATTTGGAGAATCGTTGTCAGAGCCTTACTGAGGACTTGGAGTTTCGCAAAAG
C
ATGTATGAAGAGGAGATTAACGAGACCAGAAGGAAGCATGAAACGCGCTTGGTAGAGGTGGATTCTGGG
C
GTCAAATTGAGTATGAGTACAAGCTGGCGCAAGCCCTTCATGAGATGAGAGAGCAACATGATGCCCAAG
T
GAGGCTGTATAAGGAGGAGCTGGAGCAGACTTACCATGCCAAACTTGAGAATGCCAGACTGTCATCAGA
G
ATGAATACTTCTACTGTCAACAGTGCCAGGGAAGAACTGATGGAAAGCCGCATGAGAATTGAGAGCCTT
T
CATCCCAGCTTTCTAATCTACAGAAAGAGTCTAGAGCATGTTTGGAAAGGATTCAAGAATTAGAGGACT
T
GCTTGCTAAAGAAAAAGACAACTCTCGTCGCATGCTGACAGACAAAGAGAGAGAGATGGCGGAAATAAG
G
GATCAAATGCAGCAACAGCTGAATGACTATGAACAGCTTCTTGATGTAAAGTTAGCCCTGGACATGGAA
A
TCAGTGCTTACAGGAAACTCTTAGAAGGCGAAGAAGAGAGGTTGAAGCTGTCTCCAAGCCCTTCTTCCC

G
TGTGACAGTATCCCGAGCATCCTCAAGTCGTAGTGTACGTACAACTAGAGGAAAGCGGAAGAGGGTTGA
T
GTGGAAGAATCAGAGGCGAGTAGTAGTGTTAGCATCTCTCATTCCGCCTCAGCCACTGGAAATGTTTGC
A
TCGAAGAAATTGATGTTGATGGGAAATTTATCCGCTTGAAGAACACTTCTGAACAGGATCAACCAATGG
G
AGGCTGGGAGATGATCAGAAAAATTGGAGACACATCAGTCAGTTATAAATATACCTCAAGATATGTGCT
G
AAGGCAGGCCAGACTGTTACAATTTGGGCTGCAAACGCTGGTGTCACAGCCAGCCCCCCAACTGACCTC
A
TCTGGAAGAACCAGAACTCGTGGGGCACTGGCGAAGATGTGAAGGTTATATTGAAAAATTCTCAGGGAG
A
GGAGGTTGCTCAAAGAAGTACAGTCTTTAAAACAACCATACCTGAAGAAGAGGAGGAGGAGGAAGAAGC
A
GCTGGAGTGGTTGTTGAGGAAGAACTTTTCCACCAGCAGGGAACCCCAAGAGCATCCAATAGAAGCTGT
G CAATTATGTAA

### K417-LMNB1 sequence

ATGGCGACTGCGACCCCCGTGCCGCCGCGGATGGGCAGCCGCGCTGGCGGCCCCACCACGCCGCTGAGC
C
CCACGCGCCTGTCGCGGCTCCAGGAGAAGGAGGAGCTGCGCGAGCTCAATGACCGGCTGGCGGTGTACA
T
CGACAAGGTGCGCAGCCTGGAGACGGAGAACAGCGCGCTGCAGCTGCAGGTGACGGAGCGCGAGGAGGT
G
CGCGGCCGTGAGCTCACCGGCCTCAAGGCGCTCTACGAGACCGAGCTGGCCGACGCGCGACGCGCGCTC
G
ACGACACGGCCCGCGAGCGCGCCAAGCTGCAGATCGAGCTGGGCAAGTGCAAGGCGGAACACGACCAGC
T
GCTCCTCAACTATGCTAAGAAGGAATCTGATCTTAATGGCGCCCAGATCAAGCTTCGAGAATATGAAGC
A
GCACTGAATTCGAAAGATGCAGCTCTTGCTACTGCACTTGGTGACAAAAAAAGTTTAGAGGGAGATTTG
G
AGGATCTGAAGGATCAGATTGCCCAGTTGGAAGCCTCCTTAGCTGCAGCCAAAAAACAGTTAGCAGATG
A
AACTTTACTTAAAGTAGATTTGGAGAATCGTTGTCAGAGCCTTACTGAGGACTTGGAGTTTCGCAAAAG
C
ATGTATGAAGAGGAGATTAACGAGACCAGAAGGAAGCATGAAACGCGCTTGGTAGAGGTGGATTCTGGG
C
GTCAAATTGAGTATGAGTACAAGCTGGCGCAAGCCCTTCATGAGATGAGAGAGCAACATGATGCCCAAG
T
GAGGCTGTATAAGGAGGAGCTGGAGCAGACTTACCATGCCAAACTTGAGAATGCCAGACTGTCATCAGA
G
ATGAATACTTCTACTGTCAACAGTGCCAGGGAAGAACTGATGGAAAGCCGCATGAGAATTGAGAGCCTT
T
CATCCCAGCTTTCTAATCTACAGAAAGAGTCTAGAGCATGTTTGGAAAGGATTCAAGAATTAGAGGACT
T
GCTTGCTAAAGAAAAAGACAACTCTCGTCGCATGCTGACAGACAAAGAGAGAGAGATGGCGGAAATAAG
G
GATCAAATGCAGCAACAGCTGAATGACTATGAACAGCTTCTTGATGTAAAGTTAGCCCTGGACATGGAA
A
TCAGTGCTTACAGGAAACTCTTAGAAGGCGAAGAAGAGAGGTTGAAGCTGTCTCCAAGCCCTTCTTCCC

G

TGTGACAGTATCCCGAGCATCCTCAAGTCGTAGTGTACGTACAACTAGAGGAAAGCGGGCCGAGGGTTG

AT

GTGGAAGAATCAGAGGCGAGTAGTAGTGTTAGCATCTCTCATTCCGCCTCAGCCACTGGAAATGTTTGC

A

TCGAAGAAATTGATGTTGATGGGAAATTTATCCGCTTGAAGAACACTTCTGAACAGGATCAACCAATGG

G

AGGCTGGGAGATGATCAGAAAAATTGGAGACACATCAGTCAGTTATAAATATACCTCAAGATATGTGCT

G

AAGGCAGGCCAGACTGTTACAATTTGGGCTGCAAACGCTGGTGTCACAGCCAGCCCCCCAACTGACCTC

A

TCTGGAAGAACCAGAACTCGTGGGGCACTGGCGAAGATGTGAAGGTTATATTGAAAAATTCTCAGGGAG

A

GGAGGTTGCTCAAAGAAGTACAGTCTTTAAAACAACCATACCTGAAGAAGAGGAGGAGGAGGAAGAAGC

A

GCTGGAGTGGTTGTTGAGGAAGAACTTTTCCACCAGCAGGGAACCCCAAGAGCATCCAATAGAAGCTGT

G CAATTATGTAA
