## Supplementary material for "KMT1/Suv39 methyltransferase family regulates peripheral heterochromatin tethering via histone and non-histone protein methylations": Rao and Ketkar et al_script_Computer Code 1.pdf

MATLAB Script

```
close all;
clear all;

% [FileName,PathName,FilterIndex]=uigetfile('*.lsm','Select the cell');
% fnam = strcat(PathName,FileName);
% FileName=FileName(1:end-4);s
% BW4 = imread(fnam);
% BW_cell=BW4(:,:,2);



[FileName, FilePath]=uigetfile('*.tif',...                          %No inputs, lets 
user pick a file
    'Choose tif images to import',pwd,...                           %"pwd" returns 
the folder listed as "Current Folder" in the main window
    'MultiSelect','off');                                           %Allow only one 
lsm file to be chosen
if FilePath(1)==0                                                   %If no files are 
chosen, break execution
    disp('Error in StackSlider: No files chosen');
    return
end


S.I = tiffread(strcat(FilePath,FileName));
for i=1:size(S.I,2)
    I(:,:,i) = S.I(i).data;
end

BW_cell=uint8(I(:,:,1));

figure;imshow(BW_cell,[]);

BW_cell_crop = imcrop((BW_cell),[]);
figure;imshow(BW_cell_crop,[]);

I_max= max(max(BW_cell_crop));
```

```
T = 0.1*I_max;

Im_bw_T = (BW_cell_crop > T);

% T = graythresh(BW_cell_crop);
%
% Im_bw = im2bw(BW_cell_crop,T-(0.9*T));
% figure;imshow(Im_bw,[])

se = strel('disk',1);
Im_bw = imdilate(Im_bw_T,se);
% figure;imshow(I2,[])

fh = imfill(Im_bw,'holes');

Im_L = bwlabeln(fh, 8);

S = regionprops(Im_L, 'Area');

Im_regProp = ismember(Im_L, find([S.Area] >= 2500));

figure;imshow(Im_regProp,[])

 BW1 = edge(Im_regProp,'canny');

figure;imshow(BW1,[]);

S1 = regionprops(Im_regProp,
'Centroid','MajorAxisLength','PixelIdxList','PixelList');

centroid = S1.Centroid;

% pixels =S1.PixelList;
% figure;
% Im_regProp1=flipdim(Im_regProp,1);figure;imshow(Im_regProp1,[])
figure;[C,h]=contour(Im_regProp,[1,1]);
hold on;plot(centroid(1), centroid(2),'g+');
tic
figure;imshow(BW_cell_crop,[]);
j=1;
for i = 2:size(C,2)

    xi = [C(1,i) round(centroid(1))];

    yi = [C(2,i) round(centroid(2))];

    [cx,cy,c,xi,yi] = improfile(BW_cell_crop,xi,yi,200);
```

```
    int(:,j) = c;
    j=j+1;
    if mod(i,10)==0
        hold on;plot(xi,yi,'r')
    end
end
j=0;
toc

for i=1:size(mean_int,1)
flip_plot(i,:) = mean_int(size(mean_int,1)-(i-1),:);
end

figure;plot(flip_plot);
```
